# A network model of glymphatic flow under different experimentally-motivated parametric scenarios

**DOI:** 10.1101/2021.09.23.461519

**Authors:** Jeffrey Tithof, Kimberly A. S. Boster, Peter A. R. Bork, Maiken Nedergaard, John H. Thomas, Douglas H. Kelley

**Affiliations:** Department of Mechanical Engineering, University of Rochester, Rochester, NY 14627, USA; Department of Mechanical Engineering, University of Minnesota, Minneapolis, MN 55455, USA; Center for Translational Neuromedicine, Faculty of Health and Medical Sciences, University of Copenhagen, 2200 Copenhagen, Denmark; Center for Translational Neuromedicine, Department of Neurosurgery, University of Rochester Medical Center, Rochester, NY 14642, USA

## Abstract

Rapidly growing evidence demonstrates that flow of cerebrospinal fluid (CSF) through perivascular spaces (PVSs) – annular channels surrounding vasculature in the brain – is a critically-important component of neurophysiology. CSF inflow contributes during physiological conditions to clearance of metabolic waste and in pathological situations to edema formation. However, brain-wide imaging methods cannot resolve PVSs, and high-resolution methods cannot access deep tissue or be applied to human subjects, so theoretical models provide essential insight. We model this CSF pathway as a network of hydraulic resistances, built from published parameters. A few parameters have very wide uncertainties, so we focus on the estimated limits of their feasible ranges by analyzing different parametric scenarios. We identify low-resistance PVSs and high-resistance parenchyma (brain tissue) as the scenario that best explains experimental observations. Our results point to the most important parameters that should be measured in future experiments. Extensions of our modeling may help predict stroke severity or lead to neurological disease treatments and drug delivery methods.

## Introduction

The brain lacks lymph vessels, so scientists have questioned whether a flow of cerebrospinal fluid (CSF) might play a pseudo-lymphatic role in transporting metabolic waste products (*1*). Early speculation was motivated by studies that found that tracers injected into the CSF were transported at rates faster than is possible by diffusion alone (*2, 3*). Now, renewed interest has followed the in vivo observations of Iliff et al. (*4*), who reported bulk flow of CSF through perivascular spaces (PVSs; annular channels around brain vasculature) of the murine brain (*4*), which aids clearance of amyloid-*β*, a peptide linked to Alzheimer’s disease; they named this clearance pathway the “glymphatic” (glial-lymphatic) system. Soon thereafter, Xie et al. (*5*) demonstrated that this system is active primarily during sleep. Growing evidence suggests that glymphatic dysfunction may contribute the progression of dementia (*6*) and worsened outcomes following stroke (*7*), brain trauma (*8*), and many other neurological disorders (*9*).

The glymphatic pathway is hypothesized to consist of an influx of CSF along periarterial spaces which subsequently exchanges with extracellular fluid via bulk flow, facilitated by aquaporin-4 channels on the astrocyte endfeet lining the outer wall of PVSs, followed by an efflux along perivenous spaces and nerve sheaths (*10*). Recent studies in humans have confirmed many of the key features of the glymphatic hypothesis (*11–13*). Several experimental methods have been used to probe various parts of the glymphatic system. Two-photon microscopy offers excellent temporal and spatial resolution for in vivo measurements, but typically requires invasive surgery to place a cranial window and is limited to regions near the surface of the brain (*4, 7, 14, 15*). Magnetic resonance imaging (MRI) provides noninvasive brain-wide measurements, but temporal and spatial resolution are orders of magnitude lower, rendering PVSs smaller than the spatial resolution (*11, 12, 16*). Although ex vivo analysis of brain tissue offers high resolution throughout the brain, recent studies have revealed abnormal CSF flow immediately following cardiac arrest (*17, 18*) and collapse of PVSs during tissue fixation (*15*), casting doubt on such measurements. Hence, there remains much uncertainty regarding the precise CSF flow pathway and transport rates, including glymphatic efflux routes. Resolving such details may lead to novel strategies for prevention, diagnosis, and treatment of neurological disorders (*9*).

Numerical modeling offers a powerful tool in which governing equations and physical constraints can fill voids where experimental measurements are not feasible. Indeed, much insight into the glymphatic system has already resulted from such studies (see the review articles (*19–23*)). Here we develop numerical models of CSF flow through a substantial portion of the glymphatic system and use this model to make predictions under different scenarios that account for uncertainties in important geometric and material parameters. Since a fully-resolved fluid-dynamic model is not computationally feasible, our approach employs a hydraulic network model, as in prior work (*24–27*). We investigate whether most CSF flows through the parenchyma or PVSs surrounding precapillaries, which we model as parallel pathways. Our attention to precapillary PVSs is motivated by (i) early experimental evidence of tracer transport through capillary PVSs (*3*), (ii) recent characterization of molecular markers suggesting PVSs are continuous from arterioles to capillaries to veins (*28*), and (iii) recent theoretical arguments that diffusive transport in the parenchyma coupled with advective transport in precapillary PVSs might provide an effective clearance mechanism (*20*).

In order to improve on prior idealizations of the glymphatic pathway (*25, 29*), we have developed a model of CSF flow in the murine brain based on measurements of the vascular connectivity performed by Blinder et al. (*30, 31*). We use the connectivity between different vessels in this model (Fig. 1A-C) to separately simulate either blood flow (for validation) or CSF flow. The model includes flow associated with one of the major arteries branching from the circle of Willis, e.g. the middle cerebral artery (MCA), and thus includes flow in approximately one-fifth of the cortex. MRI studies (*32, 33*) show that CSF enters pial PVSs at the circle of Willis, which is represented by the inlet node in our model, labeled in Fig. 1A-B. The model geometry for the pial vasculature (Fig. 1B) is based on a branching hexagonal model proposed in ref. (*30*), with nine pial generations amounting to 45 hexagonal units and a total of 324 penetrating arterioles. This latter value approximately matches the number of penetrating arterioles in the vicinity of the MCA, 320, which we obtained by inspecting the pial arterial reconstructions available in the Supplemental Material of Adams et al. (*34*). From data reported in ref. (*31*), we determined that, on average, 11 precapillaries branch from each of the penetrating arterioles, which we assumed to be uniformly spaced (Fig. 1C). Our hydraulic network model relates flow to the pressure differences that drive the flow and the hydraulic resistances that oppose the flow (pressure and resistance being analogous to voltage and electrical resistance in circuits). Note that this approach describes the time-averaged (net) volume flow rate and therefore neglects the oscillatory component of CSF flow, which is a reasonable approach since the Womersley number for PVS flow is small (*20*). For blood flow (or CSF flow), the resistance through the capillary bed (or capillary PVSs) and venous circulation (or venous PVSs) is modeled using single parallel resistors, shown in gray in Fig. 1C, with resistance 2.25 × 10^7^ mmHg·min/ml (or 1 mmHg·min/ml); see Methods and sections A and B in SM for details. Parenchymal flow (implemented only for CSF flow) is modeled using hydraulic resistances based on an analytical expression provided in ref. (*35*) (see C in SM). A full list of the parameters for the model is given in Table 1.

**Fig. 1:**
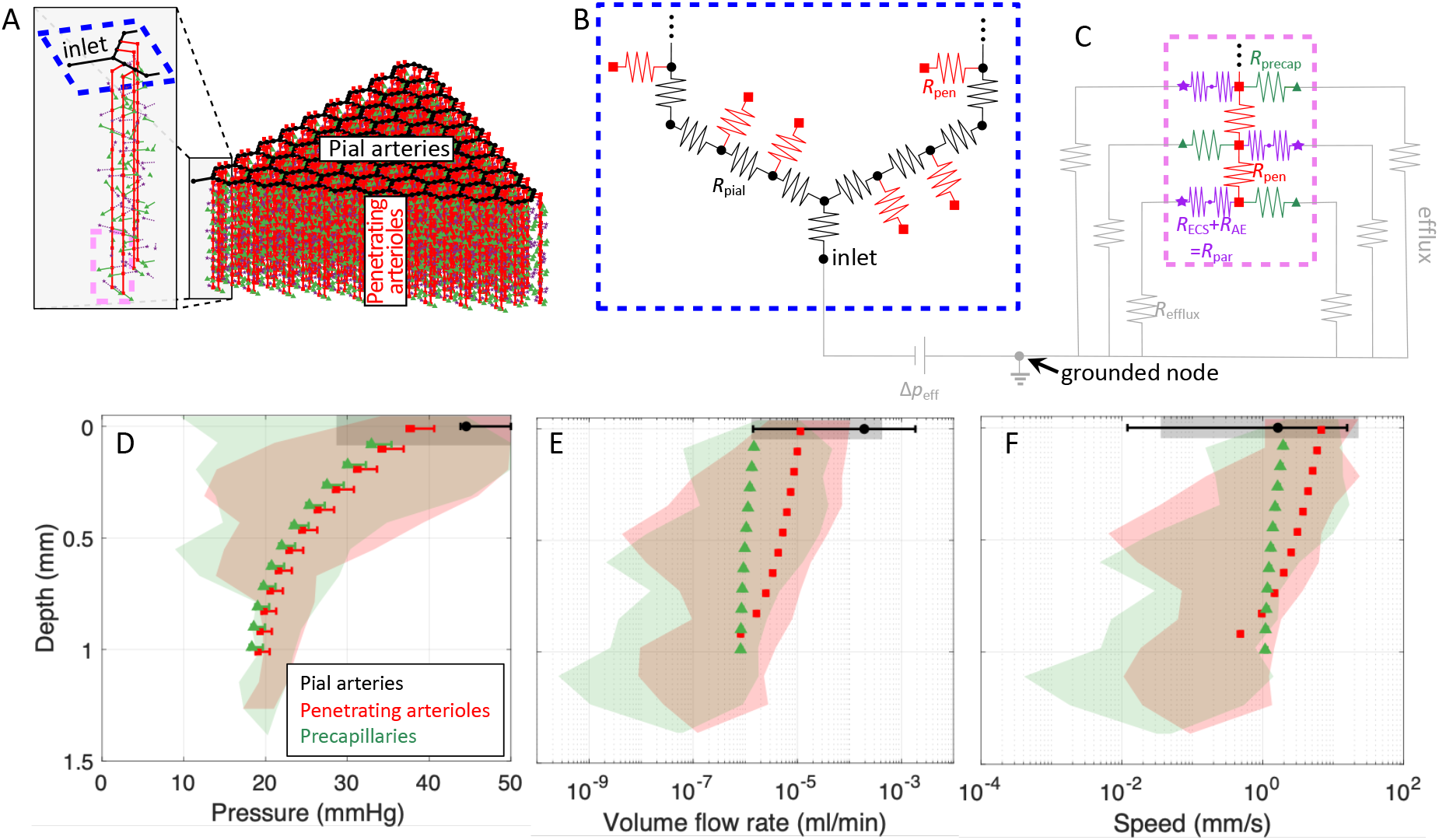
An idealized model of the cortical vasculature captures the salient features of blood flow, providing a validation of the vascular geometry used in our approach. (**A**) Diagram of the idealized vascular geometry, with colors indicating different vessel types. The blue and pink dashed lines show the regions that are enlarged in B-C. (**B**) Circuit schematic of the pial vasculature (black), which has several penetrating arterioles (red) branching from it. (**C**) Circuit schematic of a penetrating arteriole (red) which has a total of 11 precapillaries (green) branching from it (only 3 are shown). When we use a similar model to predict glymphatic CSF flow, we also include an equal number of parenchymal channels (purple). The gray circuit elements in B-C are not shown in A. (**D-F**) Pressure, volume flow rate, and speed for blood flow; in all three cases, the shaded regions indicate the range of values for a real vascular topology reported in Blinder et al. (*31*), while the symbols and error bars indicate the mean and range of values, respectively, computed using the idealized geometry shown in panel A.

**Table 1:**
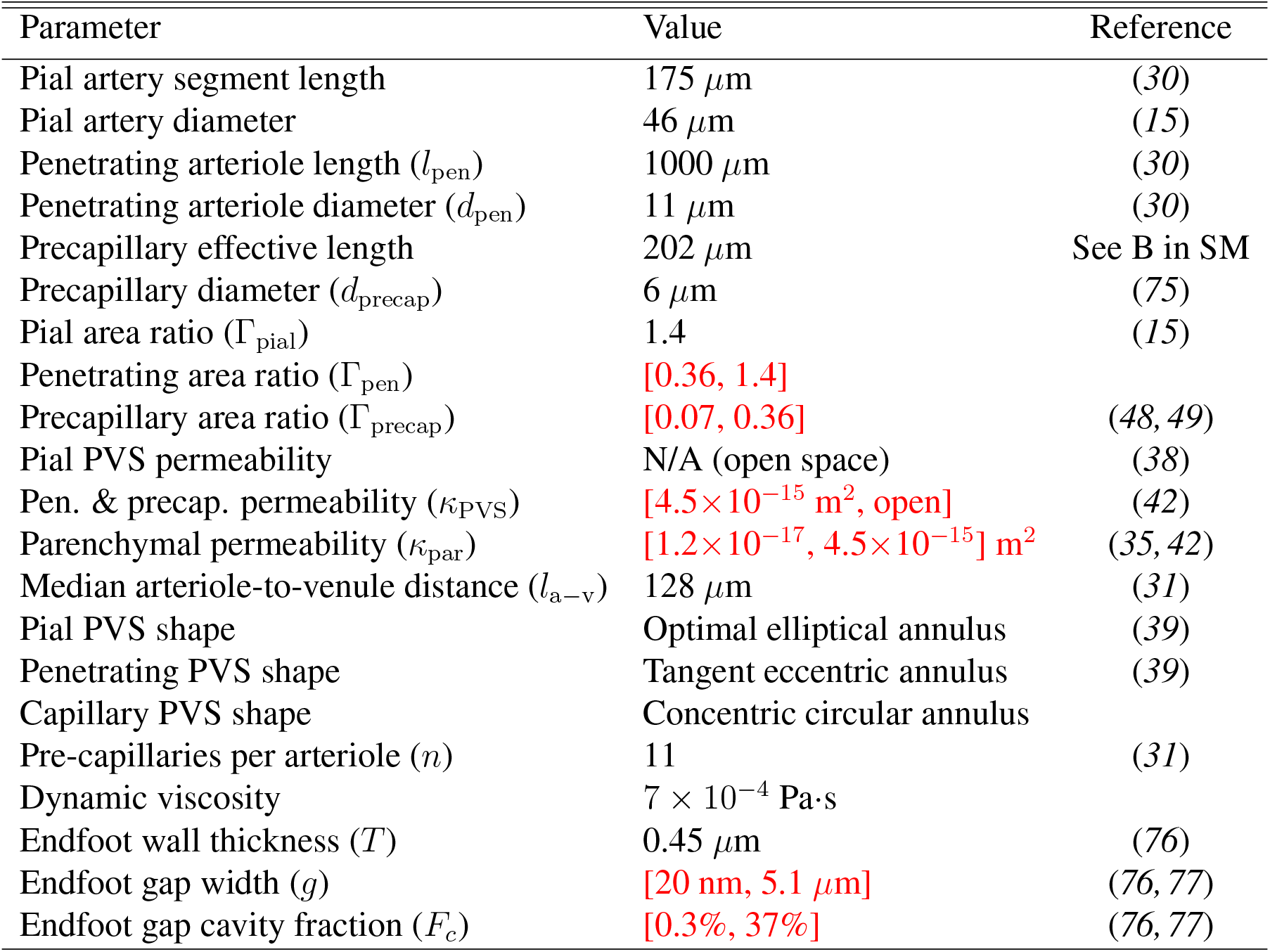
Hydraulic network model parameters. Approximate bounds for uncertain variables, which are tested in this article, are indicated in red.

| Parameter | Value | Reference |
| --- | --- | --- |
| Pial artery segment length | 175 $\mu\text{m}$ | (30) |
| Pial artery diameter | 46 $\mu\text{m}$ | (15) |
| Penetrating arteriole length ($l_{\text{pen}}$) | 1000 $\mu\text{m}$ | (30) |
| Penetrating arteriole diameter ($d_{\text{pen}}$) | 11 $\mu\text{m}$ | (30) |
| Precapillary effective length | 202 $\mu\text{m}$ | See B in SM |
| Precapillary diameter ($d_{\text{precap}}$) | 6 $\mu\text{m}$ | (75) |
| Pial area ratio ($\Gamma_{\text{pial}}$) | 1.4 | (15) |
| Penetrating area ratio ($\Gamma_{\text{pen}}$) | [0.36, 1.4] | |
| Precapillary area ratio ($\Gamma_{\text{precap}}$) | [0.07, 0.36] | (48, 49) |
| Pial PVS permeability | N/A (open space) | (38) |
| Pen. & precap. permeability ($\kappa_{\text{PVS}}$) | $[4.5 \times 10^{-15} \text{ m}^2, \text{open}]$ | (42) |
| Parenchymal permeability ($\kappa_{\text{par}}$) | $[1.2 \times 10^{-17}, 4.5 \times 10^{-15}] \text{ m}^2$ | (35, 42) |
| Median arteriole-to-venule distance ($l_{\text{a-v}}$) | 128 $\mu\text{m}$ | (31) |
| Pial PVS shape | Optimal elliptical annulus | (39) |
| Penetrating PVS shape | Tangent eccentric annulus | (39) |
| Capillary PVS shape | Concentric circular annulus |  |
| Pre-capillaries per arteriole ($n$) | 11 | (31) |
| Dynamic viscosity | $7 \times 10^{-4} \text{ Pa}\cdot\text{s}$ | |
| Endfoot wall thickness ($T$) | 0.45 $\mu\text{m}$ | (76) |
| Endfoot gap width ($g$) | [20 nm, 5.1 $\mu\text{m}$] | (76, 77) |
| Endfoot gap cavity fraction ($F_c$) | [0.3%, 37%] | (76, 77) |

## Results

### Validation of idealized network geometry via blood flow simulations

In order to validate that the idealizations of our vascular model (e.g., hexagonal connectivity, homogeneity of pial artery diameter) did not significantly alter the distribution of flow, we compared blood flow in our model with blood flow predicted for the realistic network measured by Blinder et al. (*31*). The idealized network was adjusted to cover an extent of vascular territory similar to that of the Blinder et al. study by matching the number of penetrating arterioles, resulting in a network with two pial generations (three hexagonal units), in contrast to the network shown in Fig. 1A, which consists of nine pial generations or 45 hexagonal units. In ref. (*31*), the authors measured the location and radius of all of the vessels within a section of the cortex, noting the connectivity between the vessels, and assigned a resistance to each segment based on a modified Hagen-Poiseuille law,

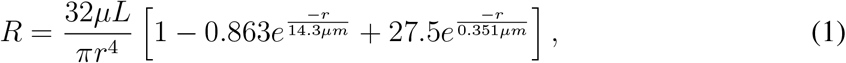

where *r* is the vessel radius, *L* is the vessel length, *R* is the resistance of that segment of vessel, and *μ* is the dynamic viscosity of water. They then applied a constant pressure difference of 50 mmHg between the arterioles and venules at the surface of the cortex and solved for the flow in each vessel. The resulting ranges of pressures, volumetric flow rates, and velocities for one mouse are indicated by the shaded regions shown in Fig. 1D-F (see Fig. S1 for results for two more mice). Based on Eqn. (1) and with a pressure difference of 50 mmHg between the inlet and outlet, we also predicted pressures, volume flow rates, and velocities for the idealized vascular geometry, which are plotted in Fig. 1D-F with solid symbols; the error bars indicate the range of values. The good agreement between the results for the realistic geometry (*31*) and for the idealized geometry indicates that the idealization does not substantially alter the salient features of blood flow through the network. The smaller range of values observed for the ide alized geometry is a result of the homogeneity of the idealization. These insights suggest that the idealized vascular geometry, which provides a framework for modeling glymphatic flow, is reasonable. Though it does not address the geometry of CSF circulation, we can infer that our results predicting glymphatic flow based on this idealized vascular geometry will likely also exhibit a narrower variation in pressure, volume flow rate, and flow speed than the actual network which has much greater heterogeneity.

### Dependence of glymphatic flow on permeability and PVS size

To model CSF flow through the glymphatic network, we enabled parenchymal flow (purple stars in Fig. 1A,C), modeled three different types of PVSs – pial, penetrating, and precapillary – and assumed homogeneity in the shapes, sizes, and porosity of each of these different PVS types (see Methods for a description of how the hydraulic resistance was computed for each pathway). Several variables needed to model fluid flow through the PVSs and parenchyma are unknown or have substantial uncertainty in their estimates. To overcome this challenge, we performed multiple simulations by bracketing the uncertain quantities (i.e., using the highest and lowest estimates of the uncertain quantities), based on a wide survey of the literature. We emphasize that in most cases, these bounds do not represent strict limits on feasible parameter ranges, but rather correspond to the extrema of values that have been reported in the literature or can be inferred from experimental data.

Bracketed parameters are indicated in red in Table 1. We considered four scenarios that lead to an overall resistance for the glymphatic network that is either maximal (*R*_max_), minimal (*R*_min_), or intermediate (Intermediate scenario 1, 2; i.e., a combination of one maximal and one minimal parameter set). For all these simulations, we matched the median pial PVS velocity to experimental measurements of 18.7 *μ*m/s (*15, 36, 37*); to obtain this match in flow speed, a different effective pressure drop Δ*p*_eff_ was required for each different scenario. We modeled the pial PVSs as open (i.e., not porous) (*38*) with a realistic, oblate shape (*39*) and a PVS-to-artery cross-sectional area ratio of Γ_pial_ = 1.4 (*15*). In vivo imaging studies suggest that pial PVSs are demarcated from the subarachnoid space (*14, 15*), although some fluid may flow between the two compartments through stomata, which are pores up to a few microns in diameter (*40*); our model does not account for stomata. For penetrating PVSs, we used either an approximate upper bound on the area ratio Γ_pial_ =1.4 (i.e., that of pial PVSs) or an approximate lower bound on the area ratio Γ_pial_ = 0.36 (i.e., the upper bound for the precapillary PVSs, discussed below). We modeled flow through the parenchyma, as well as porous penetrating and precapillary PVSs, using Darcy’s law; open (non-porous) penetrating PVSs were modeled as a tangent eccentric annulus (*39*), and open precapillary PVSs were modeled using the analytical expression for flow through a concentric annulus (see C-D in SM).

The four different scenarios we modeled arise from combining either the highest or lowest estimate of (1) the total parenchymal resistance and (2) the penetrating and precapillary PVS permeability, as detailed in Table 2. Minimum/maximum estimates of the total parenchymal resistance were obtained by lumping together the resistance from the gaps between astrocyte endfeet and the extracellular space (ECS; Fig. 2A-B; see Methods and C in SM). Note that prior studies (*24, 29, 41*) suggest CSF from penetrating PVSs primarily enters the ECS via gaps between astrocyte endfeet. The upper and lower bounds that we set on the parenchymal permeability *κ*_par_ come from two commonly cited studies (*35, 42*); multiple other studies (*43–47*) have reported *κ*_par_ values within these bounds. Basser (*42*) performed experimental measurements that estimated *κ*_par_ = 4.5 × 10^-15^ m^2^. However, Holter et al. (*35*) performed a numerical reconstruction of the neuropil, estimating *κ*_par_ = 1.2 × 10^-17^ m^2^, and speculated that the discrepancy with the earlier findings of Basser and other experimental studies may be due to fluid escaping to high-permeability pathways such as PVSs in those experiments. We therefore used this hypothesis as the basis for our R_max_ scenario, with *κ*_par_ = 1.2 × 10^-17^ m^2^ and *κ*_PVS_ = 4.5 × 10^-15^ m^2^. For the R_min_ scenario, we supposed that measurements of *κ*_par_ = 4.5 × 10^-15^ m^2^ from Basser (*42*) accurately quantify the parenchymal permeability. To model flow through penetrating and precapillary PVSs with minimal resistance, we computed an effective permeability *κ*_open_ that results from equating the volumetric flow rate predicted by Darcy’s law with the analytical expression for the volumetric flow rate for viscous flow through an open concentric circular annulus (see D in SM). This calculation defines a range of valid and invalid permeability values for a given PVS geometry, parameterized by the vessel diameter *d* and PVS-to-vessel area ratio Γ (Fig. 2C-E). We set *κ*_PVS_ equal to the value of *κ*_open_ for each corresponding geometry (penetrating and precapillary PVSs). Intermediate scenarios 1 and 2 come from choosing (1) *κ*_par_ = 1.2 × 10^-17^ m^2^ and *κ*_PVS_ = *κ*_open_ or (2) *κ*_par_ = *κ*_PVS_ = 4.5 × 10^-15^ m^2^.

**Fig. 2:**
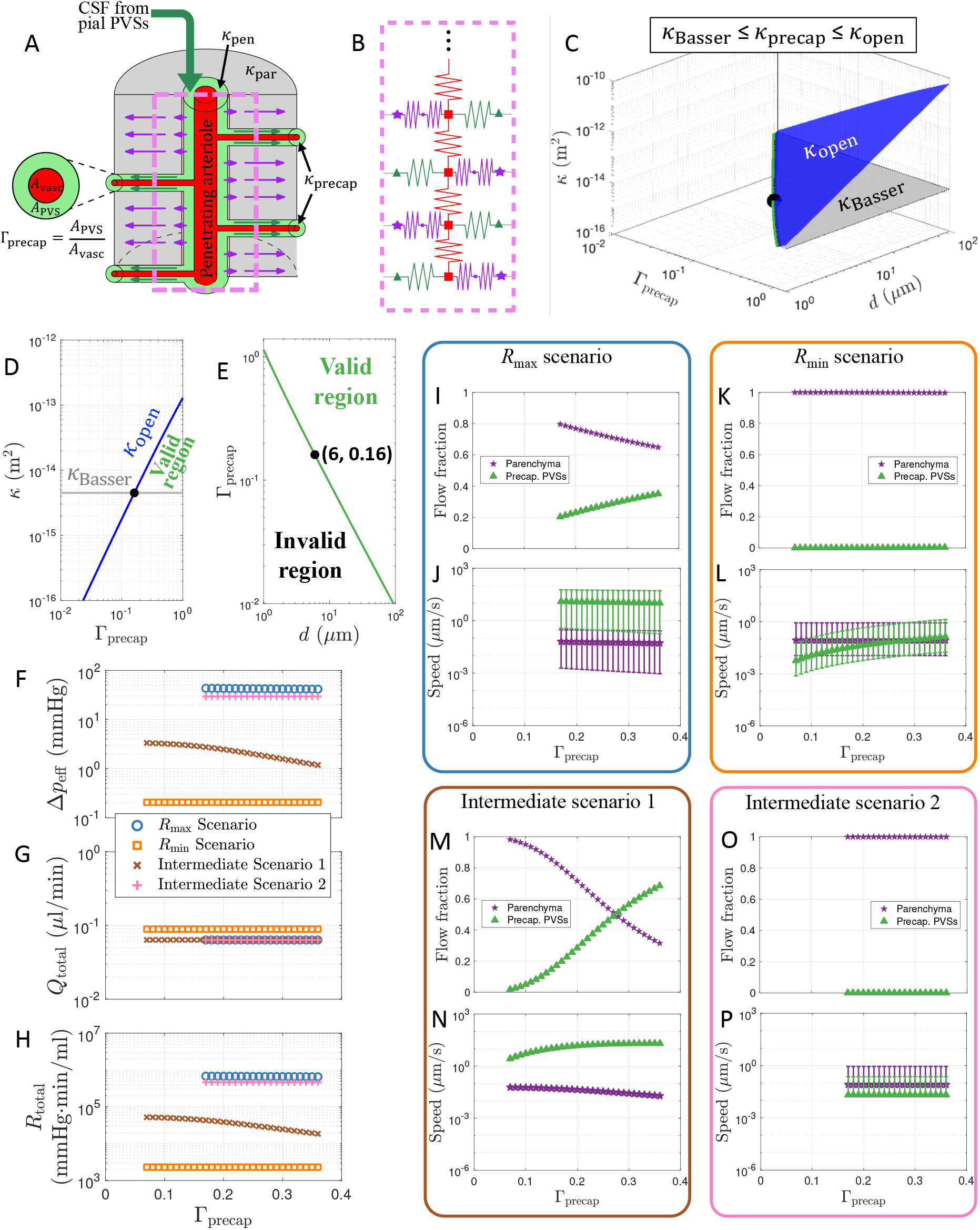
Simulations of CSF flow through the glymphatic network for different scenarios. (**A**) Schematic illustrating the geometry of a penetrating PVS segment below the cortical surface (the same segment depicted in Fig. 1C), with flow continuing through precapillary PVSs and/or the parenchyma. (**B**) Circuit schematic for the geometry shown in A (a greater portion of the network is shown in Fig. 1B-C). Throughout this article, CSF flows through the precapillary PVSs or parenchyma are consistently plotted with green or purple arrows/symbols, respectively. (**C-E**) Plots indicating the range of feasible values of permeability based on measurements performed by Basser (*42*) (*κ*_Basser_) and the equivalent permeability for an open (nonporous) PVS (*κ*_open_; see text). For d_precap_ = 6 *μ*m, PVS sizes Γ_precap_ < 0.16 are excluded for scenarios with κpvs = *κ*_Basser_ (*R*_max_ and Intermediate 2 scenarios). (**F-H**) The external pressure difference, total volumetric flow rate, and total hydraulic resistance for each of the four scenarios considered. (**I-P**) Flow fraction and flow speed through either precapillary PVSs or the parenchyma for the indicated scenarios. The symbols in panels J, L, N, and P indicate the mean flow speed across all space, while the error bars indicate the full range of values. The error bars that extend down to very low flow speeds in panels L and P arise due to negligibly small flow reaching deep into the cortex.

**Table 2:**
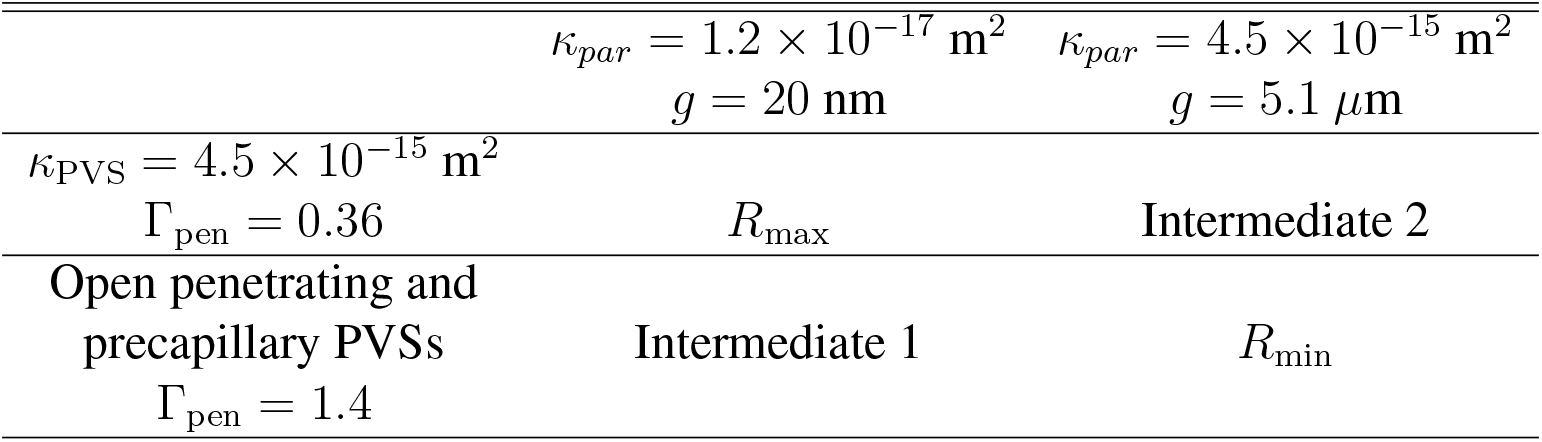
The four different parametric scenarios tested in this article. These four scenarios result from bracketing uncertain parameters related to: (left column) PVS permeability and penetrating PVS size and (top row) parenchymal permeability and astrocyte endfoot gap size.

For each of the four scenarios we considered, we varied Γ_precap_ (Fig. 2A) from 0.07 to 0.36 (i.e., the precapillary PVS gap ranged from 0.1 to 0.5 *μ*m). These values come from estimates of the size of a basement membrane (*48*) or the endothelial glycocalyx (*49*), which are the respective smallest and largest anatomical structures likely to form the contiguous portion of the PVS network at the precapillary level. In general, the anatomical details of which spaces are contiguous with penetrating PVSs are not well-understood; for a more in-depth discussion of potential PVS routes at the level of microvessels, see Hladky and Barrand (*50*). For the *R*_min_ and Intermediate 2 scenarios (with *κ*_PVS_ = *κ*_Basser_ = 4.5 × 10^-15^ m^2^), we found that an open precapillary PVS would result in an effective permeability *κ*_open_ < *κ*_Basser_ for Γ_precap_ < 0.16 (Fig. 2E). By definition, *κ*_open_ provides the upper limit on permeability, and since these two scenarios assume *κ*_Basser_ provides the lower limit of PVS permeability, we exclude Γ_precap_ < 0.16 (i.e., precapillary PVS gap widths below 0.23 *μ*m) from further analysis in these two scenarios.

The effective pressure drop Δ*p*_eff_ required to drive flow through the glymphatic network is plotted in Fig. 2F for all four scenarios. By “effective” pressure drop, we mean that we have driven flow through the network using a single pressure source (Fig. 1B), but the actual pressure gradients driving glymphatic flow – the source of which is actively debated – may be much more complex. The effective pressure drop may be thought of as *Q*_total_/*R*_total_, where *Q*_total_ and *R*_total_ are the total volume flow rate and resistance for the entire glymphatic network, respectively, even if an external pressure drop of that magnitude does not exist. Potential sources of the pressure gradients that drive the observed flows include arterial pulsations (*15*), functional hyperemia (*51*), and osmotic effects (*23, 52, 53*). The largest effective pressure drop (43 mmHg) is required for the *R*_max_ case with Γ_precap_ = 0.17, while the *R*_min_ case requires a drop of only 0.21 mmHg (which does not vary appreciably with Γ_precap_). The total volumetric flow rate through the entire network (Fig. 2G), which is approximately one-fifth of full cortical glymphatic network, varies from 0.063 to 0.089 *μ*l/min for all cases considered here; this relatively narrow range of values is a consequence of our requirement that the median flow speed in the pial PVSs match experimental measurements (*15*). With negligible variation in *Q*_total_ for a given scenario, the total hydraulic resistance of the network (Fig. 2H) is linearly proportional to the effective pressure drop, resulting in a similar functional dependence for each scenario.

We next investigated the percentage of flow that passes through the parenchyma versus the precapillary PVSs and the associated flow speed for each case (Fig. 2I-P). We found that when *κ*_par_ = 1.2×10^-17^ m^2^ (*R*_max_ and Intermediate 1 scenarios) there is a comparable fraction of total flow through the parenchyma and precapillary PVSs (Fig. 2I,M). However, if *κ*_par_ = 4.5 × 10^-15^ m^2^ (*R*_min_ and Intermediate 2 scenarios), virtually all of the flow passes through the parenchyma with a negligible amount passing through the precapillary PVSs (Fig. 2K,O). Consequently, only the former two cases show a substantial dependence on Γ_precap_, with the percentage of flow through precapillary PVSs varying from 20% to 35% as Γ_precap_ is varied from 0.17 to 0.36 for the *R*_max_ scenario, or from 1.8% to 68% as Γ_precap_ is varied from 0.07 to 0.36 for Intermediate scenario 1. The average flow speeds are plotted in Fig. 2J, L, N, P, with error bars indicating the full range of the data. The mean values for the flow speed through the parenchyma are quite similar for all four scenarios, with the average speed varying from 0.053 *μ*m/s to 0.065 *μ*m/s for the *R*_max_ scenario, or from 0.060 *μ*m/s to 0.019 *μ*m/s for Intermediate scenario 1, as Γ_precap_ is increased. For the *R*_min_ scenario (Intermediate scenario 2), the mean speed is 0.086 (0.081) *μ*m/s and does not vary appreciably with Γ_precap_. We caution that the plotted parenchymal flow speeds are not mean values across the parenchyma; they are computed at the outer wall of the PVS, so they should be interpreted as upper bounds on the parenchymal flow speed, which varies spatially. The mean precapillary flow speeds, in contrast to parenchymal speeds, show substantial variation throughout the four scenarios. The average speed varies from 13 *μ*m/s to 10 *μ*m/s for the *R*_max_ scenario, or from 2.7 *μ*m/s to 20 *μ*m/s for Intermediate scenario 1, as Γ_precap_ is increased. For the *R*_min_ scenario, the mean speed varies from 0.0058 to 0.13 *μ*m/s as Γ_precap_ is increased, but for Intermediate scenario 2 the mean speed is 0.021 *μ*m/s and does not vary with Γ_precap_. Figs. S2–S3 show how the speed and pressure vary throughout the network for the four scenarios each with maximum or minimum Γ_precap_.

### Quantifying tissue perfusion for different scenarios

Numerous studies in both humans and mice have reported that tracers injected into CSF penetrate below the brain’s surface over relatively short time scales (*11, 13, 16, 54*). Furthermore, there is growing evidence that CSF flow through the glymphatic pathway is important for the removal of metabolic waste (*13, 55, 56*), including amyloid-β (*4, 5, 57, 58*), which is produced throughout the brain. Hence, one may reasonably expect a uniform perfusion of CSF throughout the depth of the cortex to explain observations in tracer experiments and the physiological necessity of adequate waste removal. Consequently, we next computed the volume flow rate through pial PVSs, penetrating PVSs, precapillary PVSs, and the parenchyma for each of eight cases (the four scenarios introduced previously, each with either the maximum or minimum value of Γ_precap_), as shown in the left columns of each scenario in Fig. 3. It is immediately clear that when penetrating PVS resistance is minimal (*R*_min_ and Intermediate 1 scenarios), a significant volume of CSF penetrates into the deep cortex (Fig. 3E, G, I, K). However, if penetrating PVS resistance is high (*R*_max_ and Intermediate 2 scenarios), the volume flow rate drops off more rapidly with depth (Fig. 3A, C, M, O). Fig. S4 provides a visualization of how the volume flow rate varies throughout the network.

**Fig. 3:**
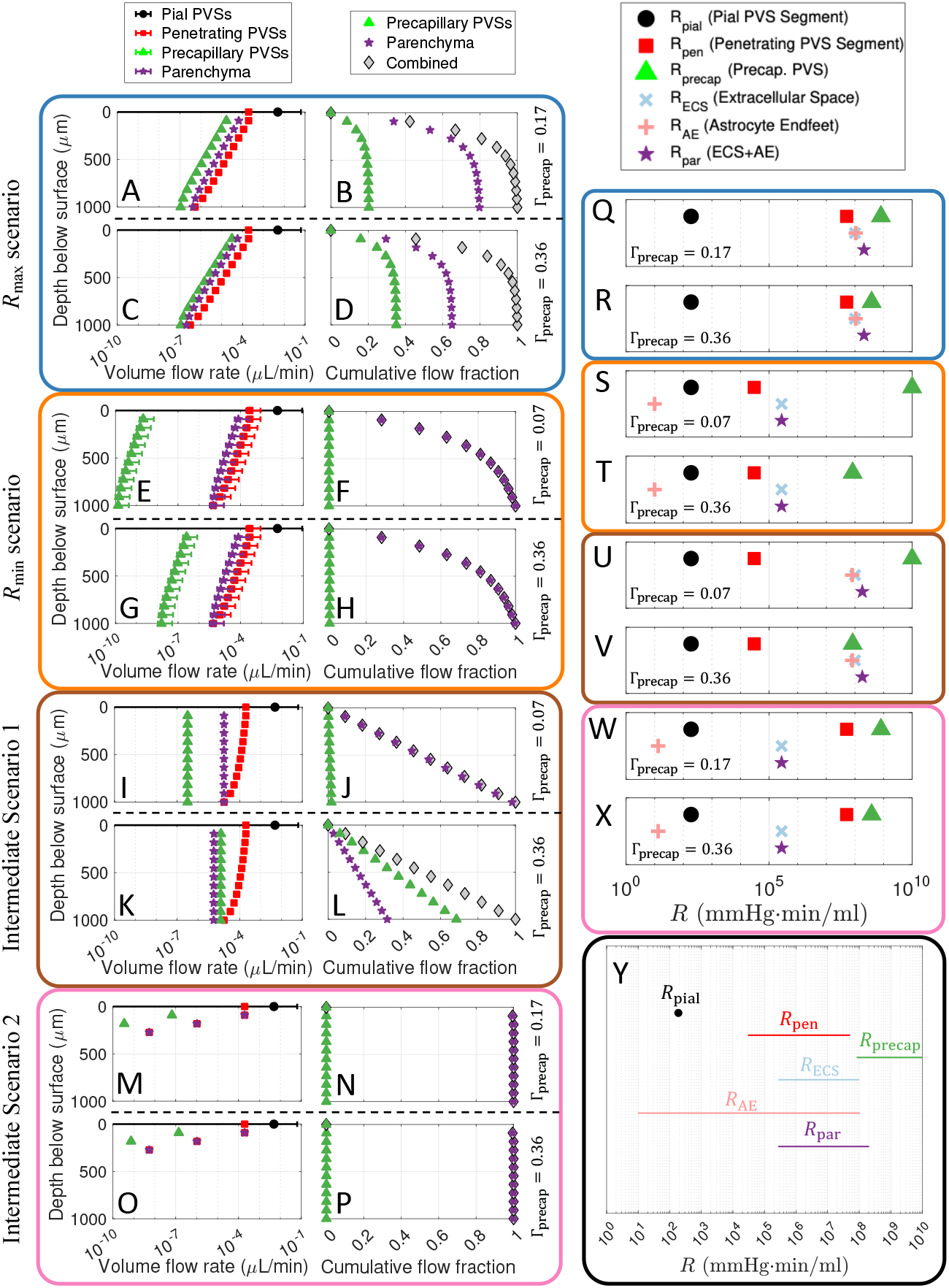
Cortical perfusion in different scenarios. (**A, C, E, G, I, K, M, O**) The volume flow rate across the depth of the cortex, and (**B, D, F, H, J, L, N, P**) the cumulative flow fraction, defined as the fraction of total volume perfused from the surface of the brain to a given depth of the cortex, for the different indicated scenarios. The legends at the top apply to each corresponding column of plots. Note that panels E-F and I-J have small precapillary PVSs (Γ_precap_ = 0.07), while panels C-D, G-H, K-L, and O-P have large precapillary PVSs (Γ_precap_ = 0.36). Panels A-B and M-N have precapillary PVSs of intermediate sizes (Γ_precap_ = 0.17) which satisfy *κ*_open_ ≥ *κ*_Basser_. (**Q-X**) Plots indicating the hydraulic resistance for a single segment of the network in each scenario, as indicated by the color of the bounding box and the Γprecap label. (**Y**) A plot of the ranges of hydraulic resistance considered across different scenarios in this study for each individual resistive element.

To characterize the perfusion, we plotted the cumulative flow fraction (i.e., the fraction of the total volume flow rate perfused from the surface of the brain to a given depth) in the right columns for each scenario in Fig. 3. The Rmax scenario has fairly poor perfusion, with 81% (Γ_precap_ = 0.17) to 84% (Γ_precap_ = 0.36) of the total CSF exiting each penetrating PVS within 270 *μ*m of the surface. Comparing the flows for small versus large precapillary PVSs (Fig. 3B,D), it is clear that more flow reroutes through the PVSs in the latter case, consistent with Fig. 2I. Among all cases, Intermediate scenario 2 exhibits the worst perfusion, with 100% of the total CSF perfusing within 270 *μ*m of the surface (Fig. 3H,P); this scenario exhibits negligible dependence on Γ_precap_. In contrast, the *R*_min_ scenario exhibits moderately uniform perfusion, with 63% of the total CSF perfusing within 270 *μ*m of the surface (Fig. 3F,N); this scenario also exhibits weak dependence on Γ_precap_. By far the best perfusion is observed in Intermediate scenario 1, for which 27% and 28% of the CSF is perfused within 270 *μ*m of the surface for Γ_precap_ = 0.07 and Γ_precap_ = 0.36, respectively (Fig. 3J, L; perfectly uniform perfusion corresponds to 27% at 270 *μ*m). Although the total perfusion remains approximately constant for different precapillary PVS sizes, as Γ_precap_ is increased a greater fraction of the flow reroutes from the parenchyma to the precapillary PVSs (compare Fig. 3I-J with K-L), consistent with the flow fractions plotted in Fig. 2M.

The variations in perfusion through the depth of the cortex for these different scenarios can be understood by comparing the hydraulic resistance of individual segments of the network, as shown in Fig. 3Q-X; several of these values are also provided in Table S1. When *R*_pen_ is substantially smaller than both *R*_par_ and *R*_precap_ (Intermediate scenario 1; Fig. 3U-V), excellent, uniform perfusion occurs (Fig. 3J, L). However, a lesser separation in resistance values (R_max_ and *R*_min_ scenarios; Fig. 3Q-T) leads to less uniformity in the perfusion (Fig. 3B,D,F,H). When *R*_pen_ is much greater than *R*_par_ (Intermediate scenario 2; Fig. 3W-X), virtually all fluid exits through the parenchymal nodes closest to the surface of the brain and perfusion is negligible at deeper nodes (Fig. 3N,P). The relative flow through the parenchyma versus precapillary PVSs can also be understood by comparing *R*_precap_ and *R*_par_. For cases where there is substantial perfusion, if the value of *R*_precap_ and *R*_par_ are comparable (Fig. 3Q-R, V), then a comparable fraction of fluid will flow through each route (Fig. 3B, D, L), with greater flow through the path of lower resistance. Two additional points are notable. The value of Rpial is much less than *R*_pen_ in every scenario, which ensures uniform perfusion of CSF across the pial PVS network (i.e., an approximately equal amount of CSF flows through both a distal penetrating PVSs and a proximal one). Also, the uncertainties in the cavity fraction and gap width of the astrocyte endfeet lead to a huge range in possible values of *R*_AE_ (Fig. 3Y). In the *R*_max_ and Intermediate 1 scenarios, the astrocyte endfeet constitute a significant barrier to flow entering the parenchyma (Fig. 3Q-R, U-V); however, in the *R*_min_ and Intermediate 2 scenarios (Fig. 3S-T, W-X), *R*_AE_ is very small and hence plays a negligible role in determining CSF flow through the parenchyma.

### Glymphatic flow during wakefulness versus sleep

We carried out additional calculations with our model aimed at investigating the increase in tracer influx during sleep/anesthesia reported by several studies (*5, 52, 59, 60*). The CSF simulations presented up to this point (Fig. 2–3) correspond to sleep conditions (or, comparably, conditions under ketamine-xylazine anesthesia). To model the change in flow during wakefulness, we used the Kozeny-Carman equation (see E in SM) to estimate that *κ*_par_ decreases by a factor of about 5.5 in wakefulness, compared to sleep. We repeated the simulations of the eight scenarios presented in Fig. 3 using a parenchymal permeability that was 5.5 times smaller, but all other parameters (including the imposed pressure drop) were left unchanged for each scenario. We then compared these results to the results from each corresponding simulation under sleep conditions. The total volume flow rates through the entire model network for wake and sleep in each of the eight scenarios are plotted in Fig. 4A-H. The combined flow for awake conditions (open gray diamonds) varies substantially across different scenarios, whereas for sleep (filled gray diamonds) all correspond to *Q*_total_ of either 0.063 or 0.089 *μ*l/min, consistent with Fig. 2G.

**Fig. 4:**
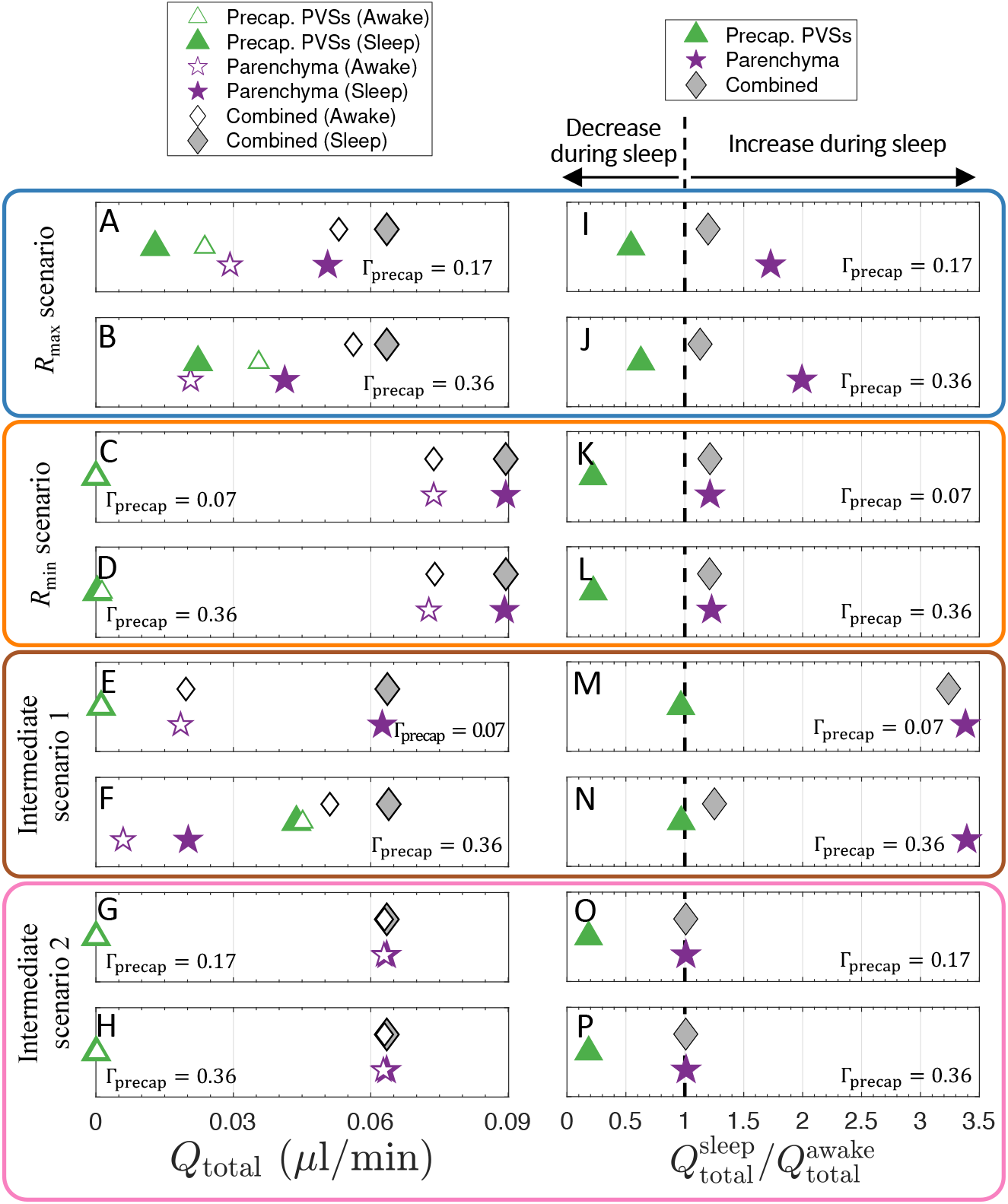
Modeled glymphatic flow in wakefulness and sleep. (**A-H**) Volumetric flow rate Qtotal summed over the entire network for different routes during either sleep or wakefulness, as indicated by the legend at the top; four different scenarios are considered, each with either small or large precapillary PVSs, as indicated. (**I-P**) The factor by which flow through precapillary PVSs, parenchyma, or both routes combined changes during sleep compared to wakefulness, quantified as Qtotal 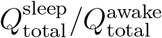 for the different indicated scenarios. The black dashed line corresponds to a value of 1, indicating no change; values to the right or left of this line correspond to an increase or a decrease, respectively, in the indicated volumetric flow rate during sleep. Note the different limiting precapillary PVS sizes Γ_precap_ indicated in the corner of each panel.

We quantified the sleep/wake change in flow by plotting the ratio of volume flow rates 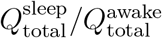, shown in Fig. 4I-P. For sleep compared to wakefulness, in every scenario the volume flow rate decreases for precapillary PVSs and increases for the parenchyma, leading to an overall increase in the combined volume flow rate. This is expected since the increased parenchymal permeability during sleep leads to an overall reduction in the hydraulic resistance of the network, and locally this change will reroute some precapillary PVS flow through the parenchyma. We find that the wake/sleep increase in combined volume flow rate is largest for small precapillary PVSs (Fig. 4I, K, M, O); this combined increase is small for Intermediate scenario 2 (0.8%), *R*_max_ (13% to 20%), and *R*_min_ (21%), but up to 220% for Intermediate scenario 1. In this latter scenario, however, there is substantial sensitivity to the size of the precapillary PVSs (Fig. 4E-F, M-N), which arises because parenchymal flow dominates the combined transport for the Γ_precap_ = 0.07 case (and is therefore sensitive to wake/sleep changes in *κ*_par_), whereas precapillary PVS flow dominates the combined transport for the Γ_precap_ = 0.36 case (and is therefore insensitive to wake/sleep changes in *κ*_par_); this observation is consistent with Fig. 2M. Of all of our wake/sleep simulations, none exhibits an increase in combined flow greater than 220%.

## Discussion

In this study, we have developed a numerical model of a substantial portion of the glymphatic system in the murine brain. This model is based on an idealized vascular geometry inspired by detailed measurements reported by Blinder et al. (*30, 31*), and we characterized the effects of idealizing the vascular geometry by first performing simulations of blood flow (Fig. 1). In modeling CSF flow through the glymphatic pathway, we matched median pial CSF velocity to experiments (*15*), we realistically modeled pial PVSs as open (nonporous) (*38*) and oblate (*39*), and we used experimentally measured mean vessel diameters and lengths. To overcome the multiple uncertainties in other parameters, we set reasonable bounds (Table 1) and performed several simulations corresponding to different combinations of the extreme values of the uncertain parameters (Fig. 2). It should be noted that these bounds are not strict extrema, but rather correspond to maximum/minimum values of each quantity as reported in various experimental studies. This “bracketing” approach included upper and lower bounds on the hydraulic resistance for penetrating PVSs, precapillary PVSs, and the parenchyma (based on a lumped model of astrocyte endfeet and the parenchymal ECS). Our model assumes CSF passes from penetrating PVSs to either precapillary PVSs or through the parenchymal ECS via a paracellular route through gaps between astrocyte endfeet (*23*). Ultimately, our goal was to investigate different scenarios to test which parameter regimes are feasible and explain as much experimental data as possible. We focused primarily on quantifying the required pressure drops, flow fraction and speed, cortical perfusion, and sleep/wake changes in volumetric flow rate.

The pressure drops and total volumetric flow rates we computed (Fig. 2F-G) provide novel insights. The two scenarios with high penetrating PVS resistance R_pen_ (R_max_ and Intermediate scenario 2) require infeasibly large pressure drops between 30 and 43 mmHg. This renders both scenarios very unlikely because such a large pressure drop is even greater than the typical systolic-diastolic variation in blood pressure of about 20 mmHg (*61*), which is thought to provide an absolute upper bound for the pressure drop driving glymphatic flow (*25*). The *R*_min_ scenario, however, requires the lowest pressure by definition, which is only 0.21 mmHg. Such a pressure is feasible and in line with estimates for the transmantle pressure difference (*62*) (i.e., that between the subarachnoid space and lateral ventricles); note however that ref. (*62*) is a model of human anatomy. For Intermediate scenario 1, the required pressure is moderately larger, varying from 1.2 to 3.3 mmHg for Γ_precap_ from 0.36 to 0.07, respectively. Such pressure is plausible, but would perhaps require driving mechanism(s) beyond simply a transmantle pressure difference (additional mechanisms are discussed further below).

Since we matched the median pial CSF velocity to experimental measurements (*15*), we find *Q*_total_ = 0.064 *μ*l/min for every scenario, except the *R*_min_ scenario in which *Q*_total_ = 0.089 (Fig. 2G). The reason *Q*_total_ is moderately larger in the the *R*_min_ scenario is because the minimal resistances of penetrating PVSs and parenchyma (Fig. 3S-T) allow more fluid to exit the network along the parenchymal nodes most proximal to the inlet, which is perhaps discernible in Fig. S4B. Our model represents approximately one-fifth of the cortical glymphatic network (e.g., in the vicinity of one MCA), so the total CSF volume flow rate through cortical PVSs would be approximately 0.32 *μ*l/min, much larger than the CSF production rate of the choroid plexus, which has recently been measured to be about 0.1 *μ*l/min for young, healthy, anesthetized mice (*63*). Although this measurement involves invasive techniques, Karimy et al. (*64*) (who developed the technique used in (*63*) in rats) reported that results were consistent with a prior method; still, this measurement may be an underestimate, as the approach excludes CSF production at the 4th ventricle.

Multiple potential explanations exist for the discrepancy between estimates of CSF production and the larger volume flow rate from our model, some of which depend on the details of pial PVSs. The pial PVSs that we have modeled are extensions of the subarachnoid space (SAS), and prior studies have suggested that not all fluid in pial PVSs continues to penetrating PVSs but rather a portion of the flow continues directly from PVSs of pial arteries to those of veins (*17, 36, 40*). Furthermore, not all CSF from the SAS enters pial PVSs; Lee et al. (*65*) delivered a tracer to the cisterna magna in rats and determined that approximately 20% reached the parenchyma, with the rest following CSF efflux routes, including the arachnoid villi, cribriform plate, and cranial and spinal nerves. Hence, it is likely that only a fraction of the total CSF enters pial PVSs, and perhaps not all CSF in pial PVSs continues through penetrating PVSs and into the parenchyma. Our prediction of a volume flow rate larger than CSF production thus suggests that either (i) published in vivo measurements of fluid velocities (*15, 36*) are inaccurately large, (ii) the pial geometry in our model is too idealized and greatly overestimates the volume flow rate, (iii) published measurements of CSF production rates are inaccurately small, and/or (iv) the fraction of CSF in pial PVSs which does not enter penetrating PVSs is able to flow back into the SAS and reenter pial PVSs of arteries, forming a kind of recirculation along the surface of the brain. Future studies could test these possibilities. Option (i) seems unlikely due to the good agreement between two independent studies (17 *μ*m/s versus 18.7 *μ*m/s reported by (*36*) and (*15*), respectively). Option (ii) perhaps plays a role, and future numerical studies with improved modeling of the pial PVS geometry should investigate this possibility. It is likely that option (iii) might contribute to the discrepancy, but such experiments are challenging and always have confounding factors. Option (iv) may contribute as well; future particle tracking experiments should investigate possibilities (i) and (iv).

The values of hydraulic resistance computed with our model can be directly compared to those of prior work. Faghih and Sharp (*25*) developed a network model of flow through periarterial spaces and computed a total network resistance of 1.14 mmHg·min/ml. This value is about 2000 times lower than the lowest hydraulic resistance we compute, R = 2300 mmHg·min/ml for the *R*_min_ scenario. This discrepancy is probably primarily because Faghih and Sharp mod eled glymphatic flow in a human, with far more parallel channels than we have considered. Vinje et al. (*27*) developed a compartmental model to estimate how elevated intracranial pressure may affect CSF outflow pathways. Although their study modeled human anatomy, they used parameters similar to Intermediate scenario 1 in this study and reported that the hydraulic resistance of the parenchyma was comparable to that of the PVSs, which is in good agreement with our observations.

We find that a substantial fraction of the CSF flowing through penetrating PVSs continues through the parenchyma in every scenario, with values ranging from 32% (Intermediate scenario 1 with Γ_precap_ = 0.36; Fig. 2M) to 100% (*R*_min_ and Intermediate 2 scenarios; Fig. 2K,O). In fact, a greater portion of CSF flows through the parenchyma than precapillary PVSs in every scenario except Intermediate scenario 1 with large precapillary PVSs (Γ_precap_ ≥ 0.27). For the *R*_max_ and Intermediate 1 scenarios, *κ*_par_ ≪ *κ*_PVS_ but in the penetrating PVSs the parenchymal-to-precapillary PVS surface area ratio is large (~ 270), leading to comparable hydraulic resistance for these two parallel pathways (Fig. 3Q-R, V). The mean parenchymal flow speeds we find are surprisingly robust across different scenarios, with values ranging from 0.019 to 0.086 *μ*m/s depending on the scenario and value of Γ_precap_ (Fig. 2J, L, N, P). Our upper bound is in agreement with the lower bound of flow speeds, 0.083 *μ*m/s, reported by Ray et al. (*66*). Additionally, our lower bound is in agreement with results from Holter et al. (*35*), in which parenchymal flow speed near the outer wall of the PVS is about 0.035 *μ*m/s (see Fig. 3 in ref. (*35*)). For cases in which the precapillary PVS flow fraction is non-negligible (> 0.5%; Rmax and Intermediate 1 scenarios), the speeds are also fairly robust, ranging from 2.7 to 20 *μ*m/s (Fig. 2J, N). This moderate insensitivity to precapillary PVS size (Γ_precap_) – especially for the *R*_max_ scenario – can be understood as follows: as the cross-sectional area A_PVS_ increases, the hydraulic resistance *R*_precap_ decreases causing the volume flow rate *Q* to increase, rendering the flow speed (= *Q/A*_PVS_) approximately constant. To the best of our knowledge, this is the first time precapillary PVS flow speed has been predicted.

We assessed whether each scenario exhibits uniformity in cortical perfusion, which we expect based on reports of tracer penetration below the brain’s surface (*11, 13, 16, 54*) and evidence that flow is important for metabolic waste removal (*4, 5, 13, 55–58*). Our simulations revealed near-perfect cortical perfusion for Intermediate scenario 1, moderately uniform perfusion for the *R*_min_ scenario, fairly poor perfusion for the Rmax scenario, and negligible perfusion below the brain surface for Intermediate scenario 2 (Fig. 3). As discussed above, good uniformity in perfusion can be understood as a consequence of scale separation in the hydraulic resistance of the three sequential CSF routes: pial PVSs, penetrating PVSs, and parenchyma/precapillary PVSs (Fig. 3Q-X). Poor perfusion occurs if these resistances are comparable (Fig. 3Q-R) or do not increase in the aforementioned order (Fig. 3W-X). This observation provides an argument in favor of large parenchymal resistance, which could arise due to tight astrocyte endfeet gaps, a low-permeability parenchymal ECS, or a combination of the two. Furthermore, the separation in scale between pial and penetrating PVS resistance ensures that CSF is uniformly perfused from the pial PVSs to the penetrating PVSs. This need for separation in scale may explain why pial PVSs have an oblate shape that minimizes their hydraulic resistance (*39*).

We performed simulations aimed at capturing the increase in CSF flow during sleep compared to wakefulness (*5, 52*). Multiple studies demonstrate that glymphatic transport is enhanced under ketamine/xylazine (K/X) anesthesia, resembling natural sleep, and inhibited under isoflurane, resembling wakefulness (*5, 33, 52, 59*); indeed, both the prevalence of slow (delta) waves and the ECS porosity under K/X are comparable to natural sleep (*5*). These studies comparing K/X and isoflurane highlight the heterogeneity of tracer transport in different regions of the brain, often with two-to four-fold greater tracer influx under K/X, compared to isoflurane. We found that *R*_min_, *R*_max_, and Intermediate scenario 2 all exhibit less than a 22% increase in combined volume flow rate during sleep compared to wakefulness (Fig. 4I-L,O-P); however, we found a 3.2-fold increase in combined volume flow rate for Intermediate scenario 1 with small precapillary PVSs (Fig. 4M). Increased tracer transport can be estimated from increased CSF flow based on the theory of Taylor dispersion (*67, 68*), which describes the effective diffusion coefficient *D*_eff_ characterizing the rate at which a tracer spreads in a shear flow due to the combined effect of advection and diffusion. For measured pial PVS size and flow speed (*15*) and a diffusion coefficient of *D* = 1 × 10^-11^ m^2^/s (*69*), *D*_eff_ /D = 3.8 (Fig. S5A), suggesting pial CSF flow enhances transport 3.8-fold greater than diffusion alone. When the awake-to-sleep volume flow rate is increased less than 22% (*R*_min_, *R*_max_, and Intermediate scenario 2), the enhanced tracer transport is less than 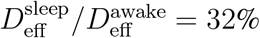, whereas a 3.2-fold increase in awake-to-sleep volume flow rate (Intermediate scenario 1 with Γ_precap_ = 0.07) leads to 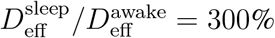 (Fig. S5B). Note that prior studies computed enhancement factors based on oscillatory (zero mean) flow (*69, 70*), whereas our calculations are based on steady (nonzero mean) flow, which we have previously argued is more effective for dispersive transport (*20, 68*). Although Taylor dispersion in pial PVSs is unlikely to account for the entirety of tracer transport observed in experiments, these estimates generally suggest that Intermediate scenario 1 with small precapillary PVSs (Γ_precap_ = 0.07) is the only scenario with sleep/awake variations in volume flow rate large enough to explain tracer transport reported in several experiments (*5, 33, 52, 59*).

Overall, we find that parameters in the general range of Intermediate scenario 1 will satisfy the majority of experimental observations described in this article. We have found that a network with low PVS resistance (high PVS permeability) and high parenchymal resistance (whether from tight gaps between astrocyte endfeet, low parenchymal permeability, or both) requires a reasonably low pressure drop (Fig. 2F), exhibits nearly perfect cortical perfusion (Fig. 3J, L), and – for small precapillary PVSs – most closely captures the observed increase in CSF influx during sleep compared to wakefulness (Fig. 4M). Additionally, Intermediate scenario 1 with Γ_precap_ ≈ 0.27 is the only case which exhibits an equal 50/50 flow through precapillary PVSs and parenchyma. It is enticing to speculate that such a parameter regime may enable dynamic regulation of CSF transport; in this scenario, if parenchymal resistance were dominated by astrocyte endfeet, small changes in the endfoot gap could substantially shift CSF perfusion between slower parenchymal flow and faster precapillary PVSs flow. We caution that the parameter space is large, so Intermediate scenario 1 does not provide the only possible case that satisfies the aforementioned criteria, but rather points to a general parametric regime.

There are numerous limitations in this study that are noteworthy. Perhaps the most consequential limitation is the uncertainty in several parameters that affect CSF transport through the glymphatic pathway, which we attempted to address by considering different limiting parametric scenarios. We restricted ourselves to a moderate number of cases for the sake of clarity, and we did so by lumping some parameters together, such as the astrocyte endfoot geometry and parenchymal permeability (Table 1). Future experimental studies aimed at refining uncertain parameters will be of tremendous value for constructing predictive models. In particular, the hydraulic resistance of gaps between astrocyte endfeet is especially uncertain, with our estimates here ranging over almost seven orders of magnitude (Fig. 3Y). The low end of this range suggests the astrocyte endfeet play no role in limiting CSF transport from the penetrating PVS to the parenchymal ECS (Fig. 3S-T, W-X), while the upper limit has hydraulic resistance comparable to that of the parenchymal ECS (Fig. 3Q-R, U-V), suggesting the astrocyte endfeet play a critical role. Indeed, a recent study (*29*) reported heterogeneity in the size of astrocyte endfeet, with larger endfeet (fewer gaps) surrounding larger vessels, which provides a mechanism that improves the uniformity of cortical perfusion. We intend to implement this feature in future simulations. In our model, CSF flow is driven by the simplest possible mechanism – an externally applied pressure drop across the entire network. However, other potential driving mechanisms (e.g., pressure gradients generated by arterial pulsations (*15*), functional hyperemia (*51*), or osmotic effects (*23, 52, 53*)) could be tested with this network model approach by implementing pressure sources (i.e., “batteries”) throughout the network. In particular, incorporation of osmotic effects could be leveraged to investigate the mechanisms by which aquaporin-4 facilitates glymphatic flow (*4, 24, 60, 71*), although there is some debate about this point (*71, 72*). Yet another important limitation to our approach, already touched on in the fourth paragraph of the Discussion, involves the connectivity of pial PVSs at the surface of the brain. By introducing “short-circuit” connections between PVSs of pial arteries and pial veins, our model could be adapted to estimate the fraction of CSF that continues along the surface of the brain versus the fraction that continues through deeper PVSs and the parenchyma. Such a model would greatly benefit from experimental estimates of how many such connections typically exist. Finally, we highlight that our model can be generalized to predict transport of dye, metabolic waste, drugs, or any other molecules due to advection-diffusion. Such future studies will contribute to the substantial ongoing debate regarding the nature of transport in penetrating PVSs (*51, 68–70*).

In future work, we intend to implement numerous refinements to our simulation, but many will likely offer improvements that are of secondary importance compared to obtaining better estimates of critical parameters (as discussed above). The idealized geometry we have adopted has a regular, repeating structure composed of four different types of homogeneous channels and consequently lacks the high spatial variability characteristic of the true network. Future models could use randomly sampled statistical distributions to assign geometric parameters (*30, 31*) or directly implement the geometry of a synthetic (*73*) or real (*7, 74*) vascular network. We restricted our model to the arterial side of the network while relying on assumptions about PVSs at the capillary and venous level to enable lumped modeling, but future studies could include substantially greater detail.

In this study, we predicted CSF transport throughout a mouse brain, but our network could be expanded to model a human brain by adding more vascular generations. Such an approach would be more challenging because of the fewer measurements available for constraining the parameter space in humans, compared to mice. However, many parameters may be conserved across species (e.g., porosity, PVS area ratios, endfoot gap size). Development of such a model has tremendous clinical value, as it could offer insight into a myriad of neurological disorders. Conditions such as Alzheimer’s disease, traumatic brain injury, and subarachnoid hemorrhage are all known to coincide with disrupted glymphatic transport (*9*).

## Materials and Methods

The network depicted in Fig. 1A was inspired by a model proposed by Blinder et al. (*30*). We used Matlab to develop the geometry, graphical representation, and computational modeling. First the spatial coordinates (for generating the schematic shown in Fig. 1A), geometry, and connectivity of the network were generated and stored. This included vessel lengths, diameters, and types (pial, penetrating, precapillary, or parenchyma). The pressures and volume flow rates throughout the network were computed by enforcing Kirchhoff’s current law, Σ*Q* = 0, at every node, where *Q* is the volumetric flow rate and summation is applied over all channels connected to a given node. To illustrate the implementation of this equation, consider three sequential nodes at pressures *p*_1_, *p*_2_, and *p*_3_ connected by channels with conductance *c*_1;2_ (which connects nodes 1 and 2) and *c*_2,3_ (which connects nodes 2 and 3). The volume flow rate from node 1 to node 2 is given by:

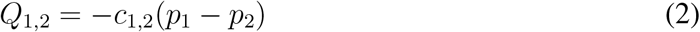

and the volume flow rate from node 2 to node 3 is given by:

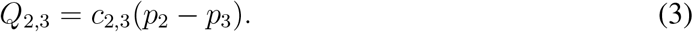

Kirchhoff’s current law requires that:

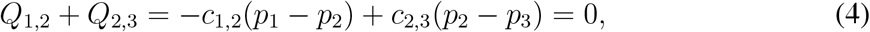

which can be rewritten as

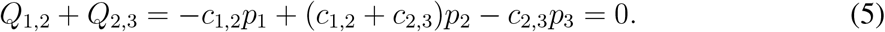

Enforcing Kirchhoff’s current law at every node in the network results in a linear algebra problem *CP* = *z* of the form:

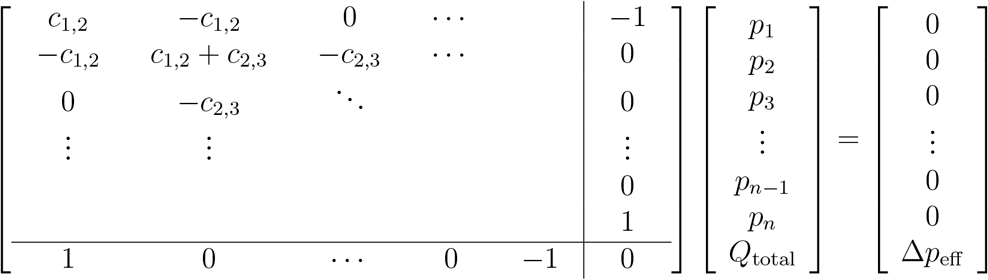

where *c_i,j_* are conductance values for the vessel connecting node *i* and *j*. Overall, the matrix *C* is sparse and was constructed by looping over each vessel segment connecting two nodes in the network and updating C with corresponding conductance values according to the connectivity of the network. Individual conductance values were computed as follows: for blood flow, Eqn. (1) was used along with a lumped model of the capillary and venous flow (see A in SM); for CSF flow, we used power laws (*39*) for nonporous pial and penetrating PVSs, the analytical solution for flow through a concentric circular annulus for nonporous precapillary PVSs (Eqn. (9) in SM), Darcy’s law for porous penetrating and precapillary PVSs, a lumped model for efflux routes (see B in SM), and another lumped model for parenchymal flow (see C in SM). The efflux node was grounded (as indicated in Fig. 1B-C) by setting the *n^th^* column of C to all zeroes. The vector P was obtained by computing the reduced row echelon form of [*C*|*z*].

Volumetric flow rates through each channel connected by nodes *i* and *j* were computed as *Q_i,j_* = *c_i,j_*(*p_i_* – *p_j_*) and the corresponding average flow speed was computed as *Q_i,j_*/*A_i,j_* where *A_i,j_* is the cross-sectional area of the given vessel or PVS (for parenchymal flow speeds, *A_i,j_* corresponds to the surface area of the outer wall of the penetrating PVS). To determine the external pressure drop Δ*p*_eff_, for a given scenario, that results in a median pial CSF flow speed of 18.7 *μ*m/s (*15*), we solved a root-finding problem. Since we want to determine the value of Δ*p*_eff_ that satisfies the equation *v*_model_(Δ*p*_eff_) = *v*_exp_, where *v*_exp_ = 18.7 *μ*m/s and *v*_model_(Δ*p*_eff_) is the median pial PVS flow speed obtained from the model, we subtract *v*_model_(Δ*p*_eff_) from both sides and define the function f (Δ*p*_eff_) = *v*_exp_ – *v*_model_(Δ*p*_eff_), which we want to equal zero. We determined Δ*p*_eff_ to an accuracy of four digits using the Matlab function “fzero”; solving this root-finding problem typically required four iterations.

Flow fractions (Fig. 2) were computed by first summing the total volumetric flow rate for either all parenchyma or all precapillary PVSs, then dividing by *Q*_total_. Cumulative flow fractions at different cortical depths (Fig. 3) were computed by summing a given volumetric flow rate (parenchyma, precapillary, or combined) for all locations at or above a given depth, then dividing by *Q*_total_. Details are provided in section E of SM describing how the change in parenchymal permeability was modeled for wakefulness relative to sleep. Total volumetric flow rates during wakefulness or sleep (Fig. 4A-H) were computed in a given scenario by summing the volumetric flow rates over the entire network for a given route (parenchyma, precapillary PVSs, or both), and each corresponding sleep/wake ratio (Fig. 4I-P) was then computed.

In addition to the validation provided by the blood flow simulations (Fig. 1D-F), we also verified our numerical methods by testing the rotational symmetry of the network, which suggests that we are indeed implementing and solving the geometry that we intend to. By implementing a total of three inlets (which is non-physiological), the network exhibits a 120° rotational symmetry (Fig. S6). We computed the relative error for each node by computing the relative error in pressure 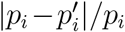, where *i* is the node index and the prime indicates the 120°-rotated network. This calculation showed that the largest deviation from rotational symmetry is 4.3 × 10^-9^%. We also verified the volumetric flow rate through each pial offshoot (i.e., the pial bifurcation leading to a penetrating PVS) by comparing Δ*p*_offshoot_/*Q*_offshoot_ to the equivalent lumped resistance for each offshoot, computed analytically. Here, Δ*p*_offshoot_ is the pressure drop between the start of each offshoot and ground and *Q*_offshoot_ is the total volumetric flow rate through a given offshoot. We find agreement in all cases to within 5.6 × 10^-5^%.

## Acknowledgments

We thank Pablo Blinder for his constructive insights and for sharing vasculature data. We also thank Dan Xue for expert assistance with Fig. S4.

## Author contributions

Conceptualization: JT, KASB, PARB, JHT, DHK

Methodology: JT, KASB, PARB

Investigation: JT, KASB

Visualization: JT

Supervision: JHT, DHK

Writing—original draft: JT, KASB

Writing-review & editing: JT, KASB, PARB, JHT, DHK, MN

## Data and materials availability

Simulation codes are available at https://doi.org/10.5281/zenodo.5644079.

## Supplementary Material

### A Lumped model of capillary bed and venous resistance for blood flow

Blinder et al. (*37*) found that the resistance across nodes in the three-dimensional resistive network of the capillary bed asymptotes to a constant value with increasing distance between nodes. They found that the asymptotic resistance is numerically the same as a network with a resistance value of 2 × 10^7^ mmHg·min/ml and that the average resistance for penetrating venules from the surface to the cortical depth layer of 4 was 2.5 × 10^6^ mmHg·min/ml. Accordingly, we used a value of 2.3 × 10^7^ mmHg·min/ml to represent the resistance to flow through the capillary bed and venous circulation back to the heart (to ground, in the circuit analogy), indicated by the gray *R*_efflux_ resistors in Fig. 1C. In this diagram, the green resistor represents the resistance to flow through a single precapillary segment, and the green symbols in Fig. 1D indicate the pressure at the distal end of that single precapillary segment.

### B Lumped model of capillary and venous PVS resistance for perivascular CSF flow

We modeled the resistance through the capillary PVSs based on the idea that the entire vascular capillary bed could be represented by a single equivalent resistor, as described by Blinder et al. (*31*). We first computed the effective precapillary length using Eqn. (1), with 5 × 10^7^ mmHg·min/ml and *r* = 2 *μ*m, consistent with the values used by Blinder et al. We used the value we obtained (202 *μ*m) to calculate the equivalent perivascular resistance. This equivalent resistance, *R*_precap_, represents the resistance to flow through the entire network of capillary PVSs beyond each given precapillary and is represented by each green resistor shown in Fig. 1C. The resistance to flow through the venous PVS, *R*_efflux_, is assumed to be negligible and is arbitrarily set as 1 mmHg·min/ml (*R*_efflux_ is represented by the gray resistors in Fig. 1C). It should be noted that this approach differs from the idealized vascular model, where *R*_precap_ represents flow through a single precapillary and *R*_efflux_ represents flow through the remainder of the capillary bed and the venous circulation.

### C Lumped model of parenchymal flow

The parenchyma was modeled as a porous medium with two-dimensional planar flow from penetrating arterioles to ascending veins. The total resistance to flow, *R*_par_, was modeled as two resistors in series, representing the resistance to flow through the gaps in the astrocyte endfeet surrounding the penetrating arteriole, *R*_AE_, and the resistance to flow through the surrounding extracellular space, *R*_ECS_, so that *R*_par_ = *R*_AE_ + *R*_ECS_. Estimates for the cavity fraction, endfeet gap width, and parenchymal permeability, which are used to calculate *R*_AE_ and *R*_ECS_ as described below, differ widely depending on the approach used to estimate them. Therefore, in order to bracket a reasonable range of expected flows, a high resistance (small cavity fraction/endfeet gap and small permeability) and a low resistance (large cavity fraction/endfeet gap and high permeability) case are modeled based on a range of estimates from the literature.

The resistance to flow through the gaps in the endfeet was modeled as flow between infinite parallel plates, for which *R* = 12*μT/g*^3^*l*, where *T* and *g* are the thickness (dimension parallel to flow) of the gap and gap width, respectively, as shown in Fig. S7. The length of the gap, *l*, was estimated by setting the area of the gap equal to the product of the cavity fraction of the gap and area of the penetrating arteriole segment through which CSF would flow, or 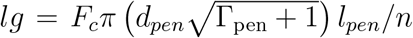, where *F_c_, d_pen_*, Γ_pen_, *l_pen_*, and *n* are the cavity fraction of the endfeet gaps, diameter of the penetrating arterioles, PVS-to-arteriole area ratio, length of the penetrating arterioles, and number of precapillaries per arteriole, respectively. Note that 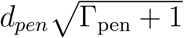 is equivalent to the diameter of the outer wall of the PVS. The resistance to flow through the endfeet gaps is then calculated as

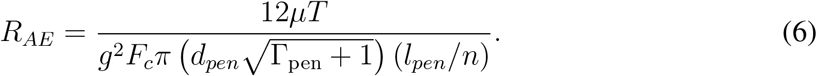

For the high resistance case, the endfoot gap and cavity fraction are assumed to be 20 nm and 0.3% based on electron microscopy measurements obtained by Mattisen et al. (*76*). Their measurements were obtained using tissue that was chemically fixed, which has been shown to significantly alter these dimensions (*77*). Nevertheless, their measurements have been used in other studies modeling the resistance to flow into the parenchyma and are included as an upper bound on the expected resistance. Korogod et al. (*77*) compared cryogenic and chemical fixation, and found significant differences in endfeet cavity fraction (37% vs 4%). For the low resistance case, we used the endfoot gap cavity fraction estimated from cryogenic fixation, 37%. Mathiisen et al. (*76*) estimated the cavity fraction they reported as *F_c_* = *gN/πd_pen_*, where *N* is the average number of transected endfoot gaps per vessel profile, which they reported as 2.5. Since the density of endfeet gaps is unlikely to change with chemical fixation, we assumed the same relationship and used *N* = 2.5 to estimate an endfoot gap width of 5.1 *μ*m for the low resistance case. In this case, *R*_AE_ was so small relative to *R_ECS_* that it could be considered negligible (Fig. 3S, T, W, X), meaning that the endfeet resist flow far less than the parenchyma.

The resistance to flow through the extracellular space, modeled as flow between a point source with constant flux to a sink, was calculated as described by Holter et al. in their Supporting Information (*35*):

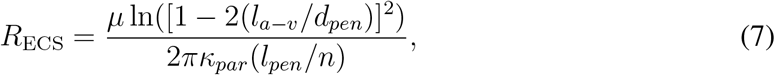

where *R*_ECS_ is the parenchymal resistance and *l_a-v_* is the median distance between an arteriole and the nearest venule. The quantity *l_pen_/n* indicates the length of the penetrating arteriole segment since the expression provided by Holter et al. was for a flux per unit length.

### D Equivalent permeability for flow through an open (non-porous) annulus

We modeled flow through the penetrating and precapillary PVSs using Darcy’s law:

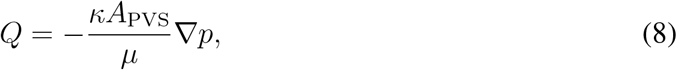

where *Q* is the volume flow rate, *κ* is the permeability, *A*_PVS_ is the PVS cross-sectional area, *μ* is the dynamics viscosity, and *p* is the pressure. To calculate the upper bound in permeability, we considered the volume flow rate through a (non-porous) concentric circular annulus, given by Eqn. (3–51) in White (*78*):

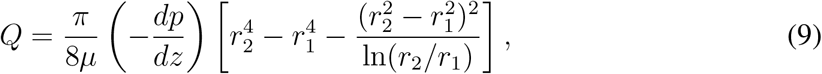

where *r*_2_ is the radius of the outer circle (outer PVS wall) and *r*_1_ is the radius of the inner circle (blood vessel). Noting that 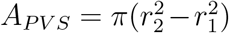, setting Eqns. (8) and (9) equal, and then solving for *κ*, one obtains:

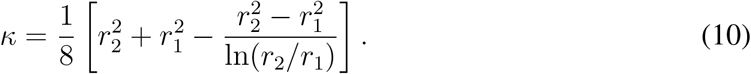

Hence, Eqn. (10) provides the upper bound for *κ*_PVS_ used throughout this article which is equivalent to modeling an open (non-porous) PVS.

### E Change in parenchymal permeability for wake versus sleep

The Kozeny-Carman equation is:

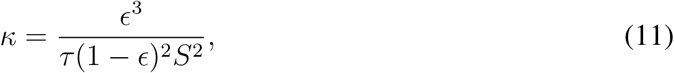

where *ϵ* is the porosity, *τ* is the tortuosity, and *S* is the specific surface area for a porous medium (*79*). Xie et al. (*5*) reported an increase of e from 0.14 during wakefulness to 0.23 during sleep, with no change in tortuosity. Assuming S remains approximately constant, this suggests 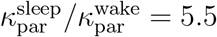.

**Table S1:** Hydraulic resistances for different circuit elements in each of the eight scenarios. The last column (*R*_offshoot_) corresponds to the entire lumped resistance for each pial offshoot, including all channels (penetrating and precapillary PVSs, parenchymal flow, and efflux) from the pial bifurcation leading to a penetrating PVS (i.e., points where the black channel bifurcates to a red channel in Fig. 1B) to ground). Units for all resistance values are mmHg·min/ml.

| Scenario | $R_{\text{pen}}$ | $R_{\text{par}}$ | $R_{\text{precap}}$ | $R_{\text{offshoot}}$ |
| --- | --- | --- | --- | --- |
| $R_{\text{max}}$ ($\Gamma_{\text{precap}} = 0.17$) | $5.131 \times 10^7$ | $1.004 \times 10^8$ | $8.114 \times 10^8$ | $1.152 \times 10^8$ |
| $R_{\text{max}}$ ($\Gamma_{\text{precap}} = 0.36$) | $5.131 \times 10^7$ | $1.004 \times 10^8$ | $3.832 \times 10^8$ | $1.116 \times 10^8$ |
| $R_{\text{min}}$ ($\Gamma_{\text{precap}} = 0.07$) | $3.018 \times 10^5$ | $2.737 \times 10^5$ | $9.986 \times 10^9$ | $1.174 \times 10^5$ |
| $R_{\text{min}}$ ($\Gamma_{\text{precap}} = 0.36$) | $3.018 \times 10^5$ | $2.737 \times 10^5$ | $8.320 \times 10^7$ | $1.173 \times 10^5$ |
| Intermediate 1 ($\Gamma_{\text{precap}} = 0.07$) | $3.018 \times 10^5$ | $1.004 \times 10^8$ | $9.986 \times 10^9$ | $9.169 \times 10^6$ |
| Intermediate 1 ($\Gamma_{\text{precap}} = 0.36$) | $3.018 \times 10^5$ | $1.004 \times 10^8$ | $8.320 \times 10^7$ | $4.271 \times 10^6$ |
| Intermediate 2 ($\Gamma_{\text{precap}} = 0.17$) | $5.131 \times 10^7$ | $2.737 \times 10^5$ | $8.114 \times 10^8$ | $6.868 \times 10^7$ |
| Intermediate 2 ($\Gamma_{\text{precap}} = 0.36$) | $5.131 \times 10^7$ | $2.737 \times 10^5$ | $3.832 \times 10^8$ | $6.868 \times 10^7$ |

**Fig. S1:**
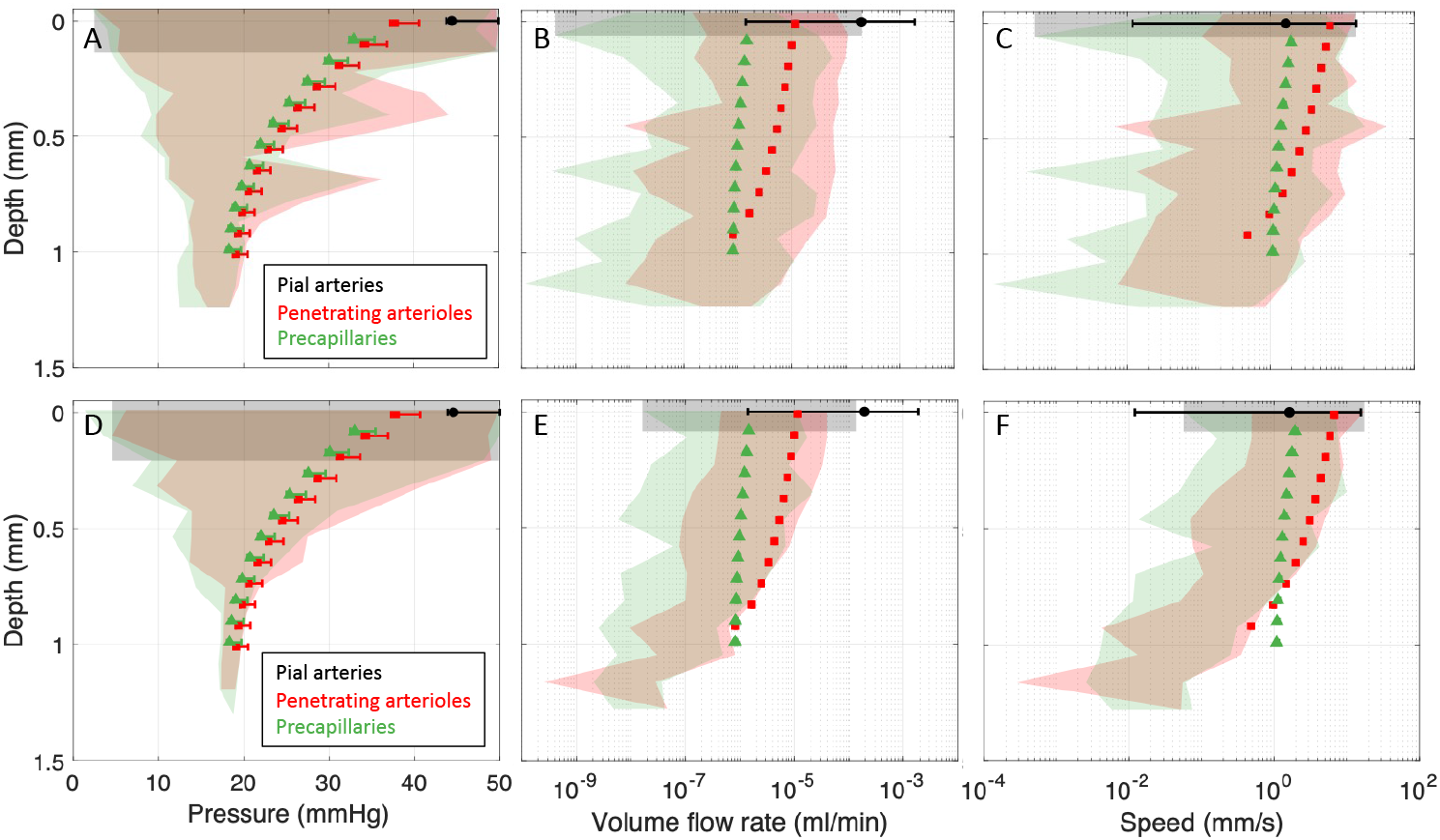
Comparisons of blood flow in the idealized model for two additional mice. Plots of (**A, D**) pressure, (**B, E**) volume flow rate, and (**C, F**) speed for blood flow in two more mice (in addition to Fig. 1D-F). The shaded regions indicate the range of values for a real vascular topology reported by Blinder et al. (*31*), while the symbols and error bars indicate the mean and range of values, respectively, computed using the idealized geometry.

**Fig. S2:**
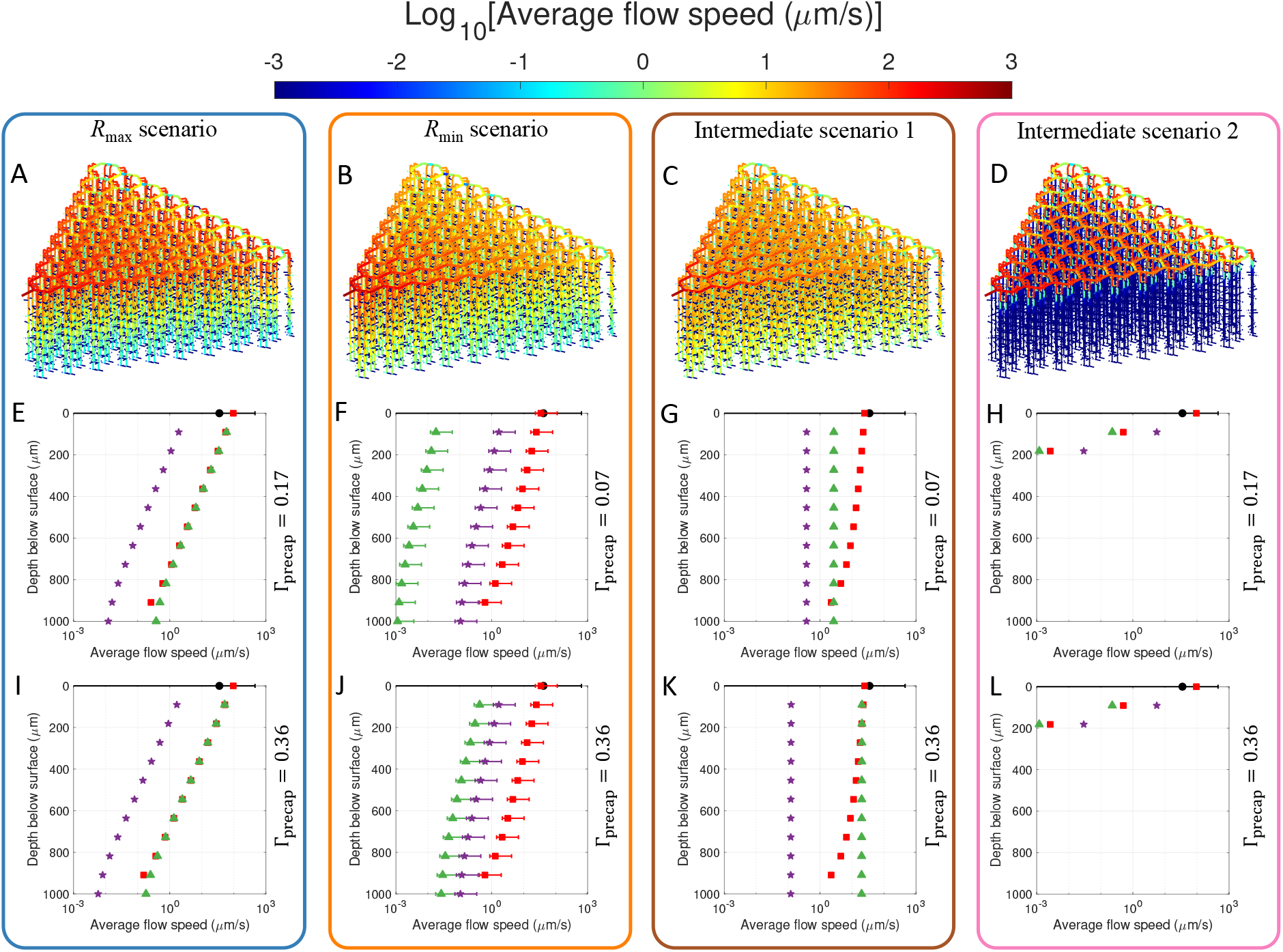
Spatial variation of flow speed. (**A-D**) Schematic diagrams of the network model with color indicating the mean flow speed. The scenarios are indicated at the top of each box; only the case of small precapillary PVSs is plotted ((A,D) Γ_precap_ = 0.17 and (B, C) Γ_precap_ = 0.07) which is virtually indistinguishable from the large precapillary PVS case. (**E-L**) Plots of average flow speed across the depth of the cortex. The error bars indicate the range of the data, and the area ratio of precapillary PVSs Γprecap is indicated to the right of each plot.

**Fig. S3:**
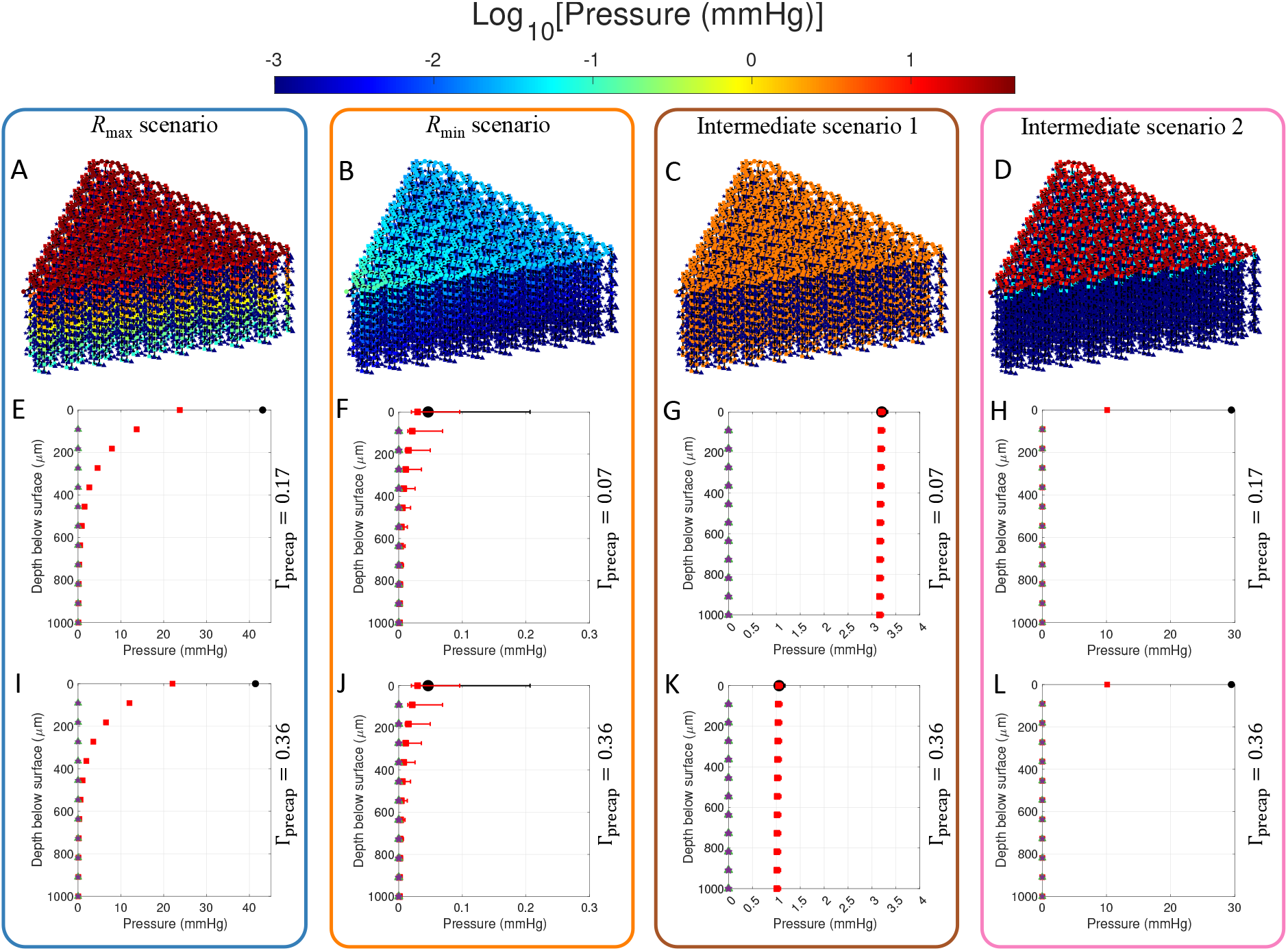
Spatial variation of pressure. (**A-D**) Schematic diagrams of the network model with color indicating the pressure, plotted on a log scale (indicated by the color bar at the top). The scenarios are indicated at the top of each box; only the case of small precapillary PVSs is plotted ((A,D) Γ_precap_ = 0.17 and (B, C) Γ_precap_ = 0.07) which is virtually indistinguishable from the large precapillary PVS case. (**E-L**) Plots of pressure across the depth of the cortex, on a linear scale. The error bars indicate the range of the data, and the area ratio of precapillary PVSs Γprecap is indicated to the right of each plot. Note the x-axis limits vary for each different scenario.

**Fig. S4:**
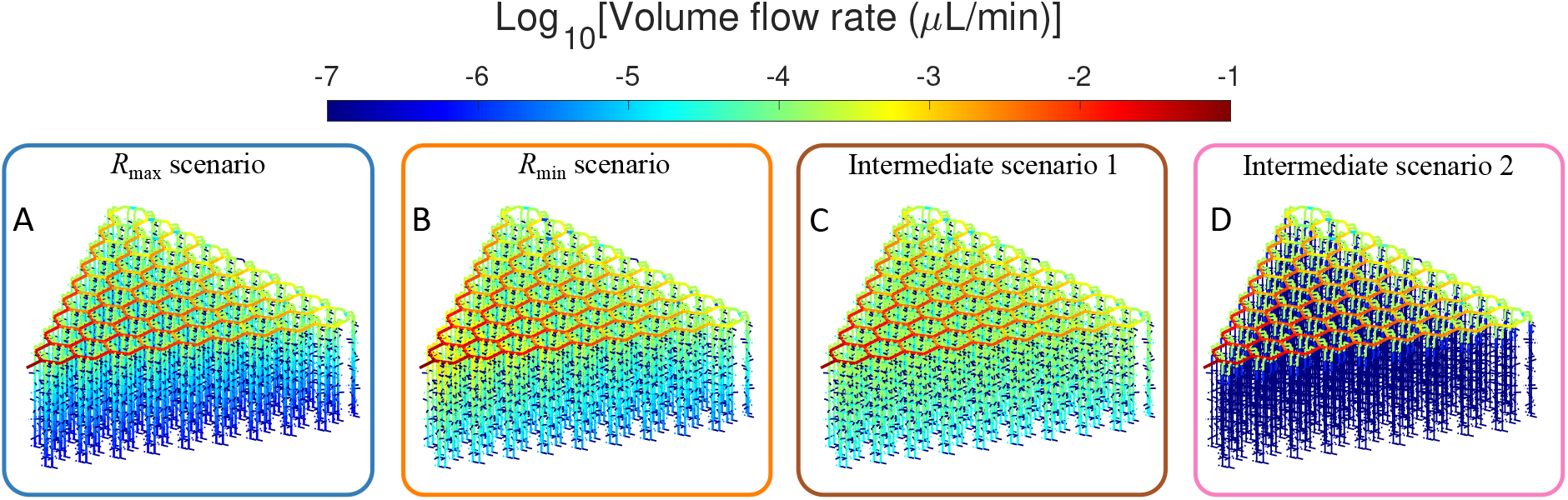
Spatial variation of volume flow rate. Schematic diagrams of the network model with color indicating the volume flow rate. The scenarios are indicated at the top of each box; only the case of small precapillary PVSs is plotted ((A,D) Γ_precap_ = 0.17 and (B, C) Γ_precap_ = 0.07) which is virtually indistinguishable from the large precapillary PVS case.

**Fig. S5:**
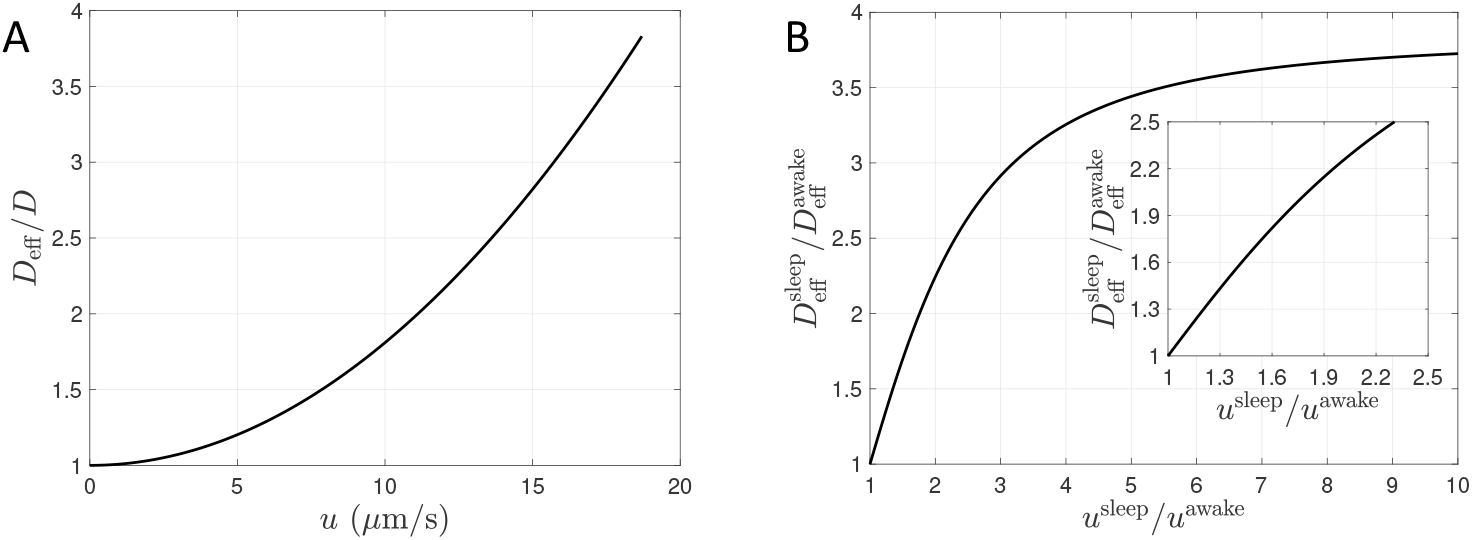
Sleep/awake dispersion coefficients for pial PVSs. (**A**) The solute transport enhancement factor *D*_eff_/*D* due to Taylor dispersion versus the mean flow speed *u*. Note that this calculation assumes the PVS shape is a concentric circular annulus with arterial diameter 46 *μ*m and Γ_pial_ = 1.4. For *u* = 18.7 *μ*m/s (*75*) and *D* = 1 × 10^-11^ m^2^/s, *D*_eff_/*D* = 3.8. (**B**) The ratio of dispersion enhancement factors 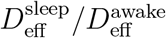 as a function of the sleep-to-awake flow speed ratio *u*^sleep^/*u*^awake^, where *u*^sleep^ = 18.7 *μ*m/s (*75*). The inset shows a magnified view for small values.

**Fig. S6:**
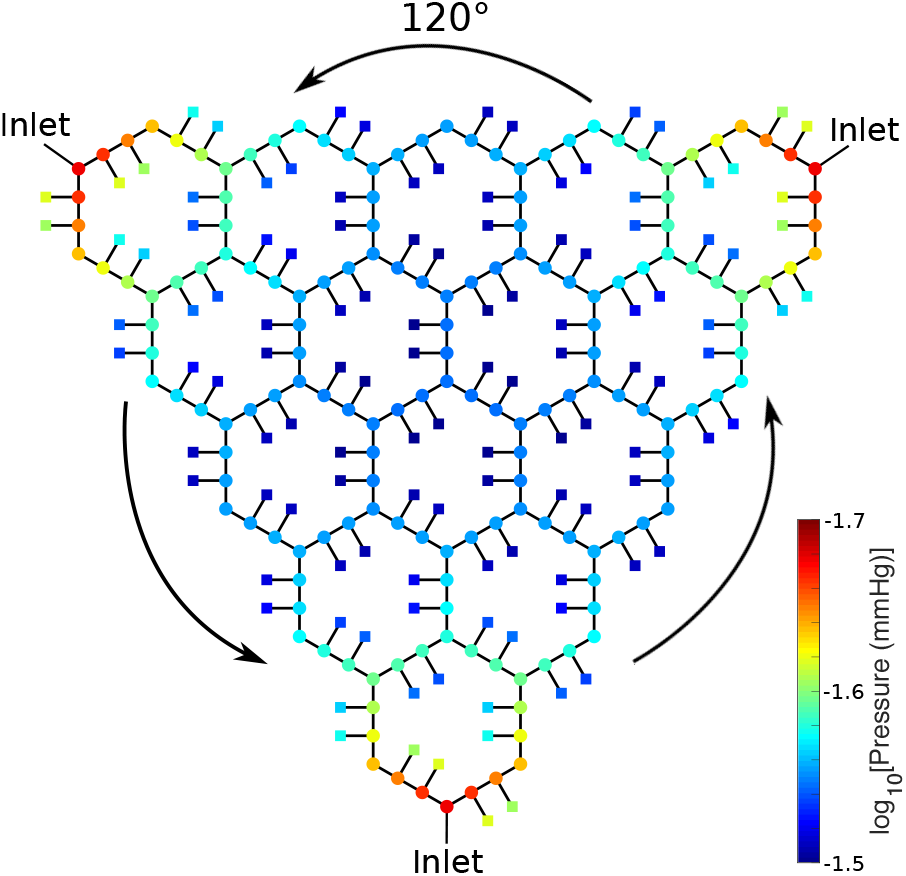
Additional model verification by testing rotational symmetry. By implementing a total of three inlets (which is non-physiological), the hydraulic network model exhibits a 120° rotational symmetry. By comparing a rotated network to the original network, we determined that the computed pressure at each node satisfies rotational symmetry to within 10^−8^%. Note that only pial nodes are plotted for the sake of clarity.

**Fig. S7:**
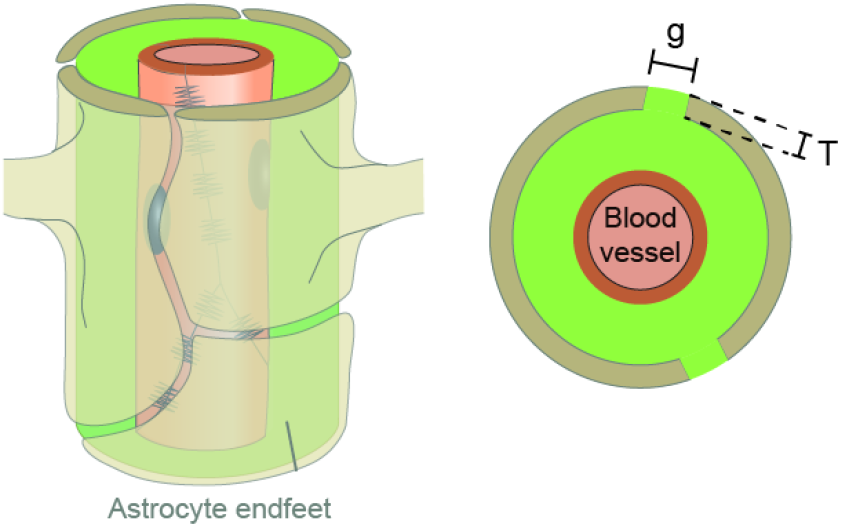
Idealized geometry of the gaps between endfeet. In our model, CSF leaves the perivascular space (green) surrounding penetrating arterioles (red) via gaps of width *g* and thickness *T*. The gaps between endfeet are long and narrow, as described by Wang et al. (*29*). To estimate the hydraulic resistance of the gaps, we consider flow between infinite parallel plates.

## References

1. T. H. Milhorat, J. Neurosurg. 42, 628 (1975).

2. H. F. Cserr, D. N. Cooper, P. K. Suri, C. S. Patlak, Am. J. Physiol.-Renal 240, F319 (1981).

3. M. L. Rennels, T. F. Gregory, O. R. Blaumanis, K. Fujimoto, P. A. Grady, Brain Res. 326, 47 (1985).

4. J. J. Iliff, et al., Sci. Transl. Med. 4, 147ra111 (2012).

5. L. Xie, et al., Science 342, 373 (2013).

6. M. Nedergaard, S. A. Goldman, Science 370, 50 (2020).

7. H. Mestre, et al., Science 367 (2020).

8. M. J. Sullan, B. M. Asken, M. S. Jaffee, S. DeKosky, R. M. Bauer, Neurosci. Biobehav. R. 84, 316 (2018).

9. M. K. Rasmussen, et al., Lancet Neurol. 17, 1016 (2018).

10. B. A. Plog, M. Nedergaard, Annu. Rev. Pathol.-Mech. 13, 379 (2018).

11. G. Ringstad, et al., JCI Insight 3 (2018).

12. N. E. Fultz, et al., Science 366, 628 (2019).

13. P. K. Eide, V. Vinje, A. H. Pripp, K.-A. Mardal, G. Ringstad, Brain 144, 863 (2021).

14. A. J. Schain, et al., J. Neurosci. (2017).

15. H. Mestre, et al., Nat. Commun. 9, 4878 (2018).

16. T. Taoka, S. Naganawa, J. Magn. Reson. Imaging 51, 11 (2020).

17. Q. Ma, et al., Acta Neuropathol. 137, 151 (2019).

18. T. Du, et al., Brain p. awab293 (2021).

19. L. A. Ray, J. J. Heys, Fluids 4, 196 (2019).

20. J. H. Thomas, J. R. Soc. Interface 16, 52 (2019).

21. H. Benveniste, et al., Gerontology 65, 106 (2019).

22. A. D. Martinac, L. E. Bilston, Biomech. Model. Mechan. 19, 781 (2020).

23. M. K. Rasmussen, H. Mestre, M. Nedergaard, Physiol. Rev. (2021).

24. M. Asgari, D. De Zélicourt, V. Kurtcuoglu, Sci. Rep. 5, 1 (2015).

25. M. M. Faghih, M. K. Sharp, Fluids Barriers CNS 15, 17 (2018).

26. J. Rey, M. Sarntinoranont, Fluids Barriers CNS 15, 20 (2018).

27. V. Vinje, A. Eklund, K.-A. Mardal, M. E. Rognes, K.-H. Støverud, Fluids Barriers CNS 17, 1 (2020).

28. M. J. Hannocks, et al., J. Cerebr. Blood F. Met. 38, 669 (2018).

29. M. X. Wang, L. Ray, K. F. Tanaka, J. J. Iliff, J. Heys, Glia 69, 715 (2021).

30. P. Blinder, A. Y. Shih, C. Rafie, D. Kleinfeld, Proc. Nat. Acad. Sci. 107, 12670 (2010).

31. P. Blinder, et al., Nat. Neurosci. 16, 889 (2013).

32. J. J. Iliff, et al., J. Neurosci. 33, 18190 (2013).

33. E. H. Stanton, et al., Magn. Reson. Med. 85, 3326 (2021).

34. M. D. Adams, A. T. Winder, P. Blinder, P. J. Drew, Sci. Rep. 8, 1 (2018).

35. K. E. Holter, et al., Proc. Nat. Acad. Sci. 114, 9894 (2017).

36. B. Bedussi, M. Almasian, J. de Vos, E. VanBavel, E. N. T. P. Bakker, J. Cerebr. Blood F. Met. pp. 0271678X1773798–8 (2017).

37. A. Raghunandan, et al., eLife 10, e65958 (2021).

38. F. Min Rivas, et al., J. R. Soc. Interface 17, 20200593 (2020).

39. J. Tithof, D. H. Kelley, H. Mestre, M. Nedergaard, J. H. Thomas, Fluids Barriers CNS 16 (2019).

40. M. E. Pizzo, et al., J. Physiol. 596, 445 (2018).

41. B. J. Jin, et al., Gen. Physiol. 148, 489 (2016).

42. P. J. Basser, Microvasc. Res. 44, 143 (1992).

43. P. F. Morrison, D. W. Laske, H. Bobo, E. H. Oldfield, R. L. Dedrick, Am. J. Physiol. - Reg. I. 266, R292 (1994).

44. S. S. Prabhu, et al., Surg. Neurol. 50, 367 (1998).

45. R. H. Bobo, et al., Proc. Natl. Acad. Sci. 91, 2076 (1994).

46. K. B. Neeves, C. T. Lo, C. P. Foley, W. M. Saltzman, W. L. Olbricht, J. Control. Release 111, 252 (2006).

47. J. H. Smith, J. A. C. Humphrey, Microvasc. Res. 73, 58 (2007).

48. P. D. Yurchenco, CSH Perspect. Biol. 3, a004911 (2011).

49. S. Reitsma, D. W. Slaaf, H. Vink, M. A. M. J. Van Zandvoort, M. G. A. oude Egbrink, Pflug. Arch. Eur. J. Phy. 454, 345 (2007).

50. S. B. Hladky, M. A. Barrand, Fluids Barriers CNS 15, 30 (2018).

51. R. T. Kedarasetti, et al., Fluids Barriers CNS 17, 1 (2020).

52. B. A. Plog, et al., JCI insight 3 (2018).

53. G. Halnes, K. H. Pettersen, L. Øyehaug, M. E. Rognes, G. T. Einevoll, Computational Glioscience (Springer, 2019), pp. 363–391.

54. T. Gaberel, et al., Stroke 45, 3092 (2014).

55. S. Koundal, et al., Sci. Rep. 10, 1 (2020).

56. X. Gu, et al., J. Control. Release 322, 31 (2020).

57. K. F. Roberts, et al., Ann. Neurol. 76, 837 (2014).

58. E. Shokri-Kojori, et al., Proc. Natl. Acad. Sci. 115, 4483 (2018).

59. L. M. Hablitz, et al., Science Adv. 5, eaav5447 (2019).

60. L. M. Hablitz, et al., Nat. Commun. 11, 1 (2020).

61. D. L. Mattson, Am. J. Hypertens. 14, 405 (2001).

62. R. D. Penn, A. Linninger, Pediatr. Neurosurg. 45, 161 (2009).

63. G. Liu, et al., Cell Rep. 33, 108524 (2020).

64. J. K. Karimy, et al., J. Neurosci. Meth. 241, 78 (2015).

65. H. Lee, et al., Magn. Reson. Med. 79, 1568 (2018).

66. L. Ray, J. J. Iliff, J. J. Heys, Fluids Barriers CNS 16, 6 (2019).

67. G. I. Taylor, P. R. Soc. A 219, 186 (1953).

68. D. E. Troyetsky, J. Tithof, J. H. Thomas, D. H. Kelley, Sci. Rep. 11, 4595 (2021).

69. M. Asgari, D. de Zélicourt,, V. Kurtcuoglu, Sci. Rep. pp. 1–11 (2016).

70. M. K. Sharp, R. O. Carare, B. A. Martin, Fluids Barriers CNS 16, 13 (2019).

71. H. Mestre, et al., eLife 7, e40070 (2018).

72. A. J. Smith, X. Yao, J. A. Dix, B.-J. Jin, A. S. Verkman, eLife 6, e27679 (2017).

73. A. Linninger, G. Hartung, S. Badr, R. Morley, Comput. Biol. Med. 110, 265 (2019).

74. C. Kirst, et al., Cell 180, 780 (2020).

75. T. Miyawaki, et al., Nat. Commun. 11, 1 (2020).

76. T. M. Mathiisen, K. P. Lehre, N. C. Danbolt, O. P. Ottersen, Glia 58, 1094 (2010).

77. N. Korogod, C. C. H. Petersen, G. W. Knott, eLife 4, e05793 (2015).

78. F. M. White, Viscous Fluid Flow (McGraw-Hill, New York, 2006), third edn.

79. J. Hommel, E. Coltman, H. Class, Transp. Porous Med. 124, 589 (2018).

